# Decoding Allosteric Grammar with Explainable AI Integrating Protein Language Models and Energy Landscape Analysis: Neutral Frustration at Allosteric Binding Sites Encodes Regulatory Versatility in Protein Kinases

**DOI:** 10.64898/2026.03.21.713412

**Authors:** Will Gatlin, Max Ludwick, Lucas Turano, Brandon Foley, Kamila Riedlová, Vít Škrhák, Marian Novotný, David Hoksza, Gennady M. Verkhivker

## Abstract

Allosteric regulation enables protein kinases to integrate diverse cellular signals, yet the energetic organization principles that encode spatial and evolutionary diversity of regulatory binding sites lack a complete understanding. We introduce an explainable artificial intelligence (AI) framework that uses protein language models (PLMs) not as predictive endpoints, but as diagnostic probes of biophysical organization of regulatory regions. By integrating PLM-based binding site predictions with the energy landscape-based frustration analysis, we determine that the detectability of orthosteric and allosteric binding sites reflects their energetic embedding within the protein energy landscape. Using PLM predictions as unbiased probes across a structurally and functionally diverse dataset of 453 human kinases, we observe a striking and reproducible dichotomy in predictive behavior: orthosteric ATP-binding pockets are detected with high confidence, whereas allosteric sites consistently evade robust detection. Orthosteric catalytic sites reside within minimally frustrated, optimized energetic regions that are consistently detected with high confidence. In contrast, allosteric sites are enriched in neutrally frustrated zones, producing diffuse and context-dependent predictions, revealing that the “allosteric blind spot” arises from intrinsic biophysical design rather than algorithmic limitations. Atomic-resolution analysis of ABL kinase spanning multiple conformational states and complexes bound to diverse ligands provides mechanistic validation of this principle. The myristoyl allosteric pocket in ABL remains neutrally frustrated across complexes with physiological ligands, chemically diverse modulators, from allosteric inhibitors to activators, and conformations engaged with SH2–SH3 regulatory domains. We propose that allosteric sites are encoded in persistent neutrally frustrated regions optimized for context-dependent regulatory modulation. By using explainable AI to interrogate the energetic architecture of protein kinases, this work reveals how the organization of the protein energy landscape shapes functional plasticity and algorithmic detectability of regulatory binding sites.

## Introduction

Allosteric interactions and communications in proteins are central to diverse biological mechanisms, and allosteric regulation has long been recognized as the “second secret of life”.^1–7^ Nonetheless, the diverse dynamic mechanisms that give rise to allosteric events continue to be fairly elusive and are often characterized at a phenomenological level, lacking a universal theoretical foundation that can explain the enormous diversity of these processes. This gap between experimental observation and mechanistic understanding is particularly acute at the intersection of allosteric regulation and the rapidly advancing fields of machine learning (ML) and artificial intelligence (AI) -a frontier where AI-augmented integrative structural biology holds the promise of transforming how we discover and interpret the regulatory logic of allostery encoded in protein structures.^8,9^ Recent years have witnessed the remarkable growth of data-intensive experimental high-throughput technologies and AI/ML tools aiming for atomistic-level interrogation, characterization and engineering of protein functions and mechanisms. Of particular interest for rapidly evolving AI tools in biology is accurate prediction of diverse regulatory allosteric sites in protein structures and dissecting biophysical principes that govern how certain regions of proteins emerge hubs of regulatory control while others remain structurally inert.

In this work, we approach this longstanding question from a new perspective: rather than using AI solely as a predictive tool, we introduce and employ an explainable AI framework as a diagnostic instrument to interrogate the global energetic organization of proteins and embedding of regulatory regions, asking whether the predictive behavior of modern AI tools and particularly PLMs can reveal the physical principles that encode diversity of allosteric regulatory sites within protein structures.

Recent advances in large-scale mutational profiling have begun to illuminate the global architecture of allosteric regulation. Pioneering studies from Lehner and colleagues used deep mutational scanning (DMS) and thermodynamic modeling to systematically map the energetic consequences of mutations across protein interaction domains.^10–13^ These experiments revealed that allostery is not confined to a small set of specialized residues but instead emerges from distributed networks of interactions spanning entire protein domains. Importantly, these studies showed that allosteric landscapes exhibit a modular organization in which conserved structural cores are surrounded by more variable peripheral regions that diversify across homologs. They proposed a generalized mechanism where allostery evolves through the gain and loss of peripheral extensions to this conserved core, allowing for functional diversification while maintaining a fundamental allosteric mechanism.^13^ These illuminating findings suggest that regulatory potential may be encoded not simply in individual residues but in the broader energetic organization of the protein structure and the underlying energy landscape where rigid protein cores communicate with flexible regions to mediate context-dependent regulation -a principle that directly resonates with the frustration-based explainable AI framework we develop in the current investigation.

Protein kinases provide a particularly powerful system for exploring these ideas. Allosteric regulation has long been viewed as the hidden language of cellular signaling^14^ and yet biophysical grammar that encodes regulatory diversity at spatial and evolutionary level remains elusive. As central regulators of cellular signaling, protein kinases are by far among the best-characterized protein families both structurally and functionally, with thousands of experimental structures spanning multiple conformational states and ligand-bound forms.^15,16^ The extensive structural and biochemical studies have revealed how functional elements of the kinase domains reorganize during activation and inhibition, producing a rich mechanistic picture of kinase regulation.^17–19^ This wealth of structure-functional data, combined with rigorous classification systems that distinguish orthosteric ATP-competitive inhibitors from proximal and distal allosteric modulators^20–22^ enables systematic comparison of different classes of binding sites and regulatory region within a common structural framework. Moreover, protein kinases possess a remarkable diversity of regulatory interfaces distributed across their catalytic domains that reflects an evolutionary strategy in which signaling enzymes exploit multiple structural sites to fine-tune activity and integrate diverse cellular signals, providing a rich landscape for investigating how allosteric sites are encoded.^23–26^

While experimental studies have revealed the richness of allosteric regulation, computational prediction of regulatory regions and particularly allosteric sites remains a major challenge. Deep learning (DL) approaches and PLMs trained on large protein sequence datasets^27–31^ have achieved remarkable success in in proteome-wide functional site annotation, with particular strength in identifying conserved binding sites.^32–35^ Structure-based ML algorithms^36–39^ exploited geometric and physicochemical features to robustly identify canonical orthosteric ligand-binding pockets in protein structures. Despite these advances, robust detection of functional allosteric sites remains challenging, as most methods are optimized for stable, evolutionarily conserved canonical binding pockets, while allosteric sites are often transient, weakly conserved, and sparsely represented in structural databases.^40^ A recent study combined local binding geometry, coevolutionary information, and dynamic allostery into a multiparameter framework designed to identify potentially hidden allosteric sites within ensembles of protein structures bound to orthosteric ligands.^41^ The emerging dichotomy in predictions of canonical orthosteric and regulatory allosteric binding sites is commonly attributed to limited training data or methodological inefficiencies. While these factors undoubtedly contribute to a markedly reduced prediction performance of PLMs and ML tools on allosteric sites, a more granular analysis from experimental studies suggests a deeper explanation. If regulatory sites occupy regions that are evolutionarily permissive and structurally adaptable-features necessary for accommodating diverse regulatory interactions-then they may lack the strong sequence conservation patterns that machine learning models rely upon. In this view, the apparent “blind spot” of AI algorithms may reflect the intrinsic evolutionary design of regulatory regions rather than a deficiency of current predictive methods.

Energy landscape theory provides a powerful framework for exploring this possibility.^42,43^ Rather than viewing proteins as static structures, this perspective describes them as ensembles of conformational states organized on a multidimensional energy landscape whose topology governs folding, dynamics, and functional transitions. Local frustration analysis offers a quantitative approach for interrogating this landscape by evaluating how favorably individual interactions are optimized relative to alternative configurations.^42–44^ Evolution tends to minimize frustration in regions essential for structural stability, creating strongly funneled energetic basins that ensure robust folding. In contrast, regions involved in binding, signaling, or conformational switching often retain higher levels of frustration, enabling conformational flexibility and facilitating functional transitions.^44^ Across many protein families, minimally frustrated residues cluster in conserved structural cores, whereas frustrated or neutrally frustrated interactions often appear at ligand-binding pockets, protein–protein interfaces, and conformational switch regions.^44–47^ The frustration patterns have been linked to evolutionary conservation, conformational dynamics, and allosteric communication pathways.^45–47^ Combined, these observations raise the possibility that the energetic embedding of binding pockets within the protein landscape may determine both their regulatory behavior and their evolutionary signatures.

In this study, we integrate PLM predictions of orthosteric and allosteric binding sites with the energy landscape analysis into an explainable dual AI framework to probe protein allostery and interrogate the organization of regulatory sites in protein kinases.. Rather than using PLMs solely as a predictive tool, we employ a physics-centric explainable AI as a diagnostic instrument and biophysical lens to probe the energetic embeddings of diverse binding sites and regulatory regions in the global architecture of protein kinase structures. Our investigation unfolds in three integrated steps. First, we systematically characterize the differential detectability of orthosteric and experimentally characterized allosteric sites using PLM predictions across a curated dataset of 453 human kinases from the KinCoRe database.^21^ This analysis establishes that orthosteric and allosteric sites occupy distinct regimes of algorithmic visibility-a pattern that raises questions about interplay and balance between methodological limitations and biophysical origins. Second, we introduce local frustration analysis as a biophysical interpretability layer, testing whether the observed performance patterns correspond to underlying differences in how these sites are embedded within the energy landscape. By comparing frustration signatures across site classes and mapping their spatial distribution, we ask whether orthosteric and allosteric sites exhibit fundamentally different energetic organization. Third, we examine these principles at atomic resolution using the ABL kinase system, that provides a uniquely well-characterized example of divers allosteric regulatory hubs. By analyzing a comprehensive set of ABL structures-including complexes with orthosteric inhibitors, allosteric modulators of opposing function, and assemblies with regulatory domains-we investigate how conformational and mutational frustration patterns manifest across different regulatory contexts and examine how neutral frustration persists across conformational states and enables functional versatility of the diverse allosteric sites in human kinome. Our results reveal a consistent organizing principle: orthosteric catalytic sites are embedded within minimally frustrated regions of the energy landscape that are strongly constrained by evolution and therefore readily detected by AI models, whereas allosteric sites are encoded within neutrally frustrated zones of the energy landscape which underlies their regulatory versatility and limited detectability by AI models. More broadly, these findings demonstrate how explainable AI can be used not only to predict functional regions and binding sites, but to serve as diagnostic probes of the physical principles governing their evolutionary design, transforming AI tools from a predictive technology into a powerful instrument for interrogating protein function.

## Results

### An Explainable AI Framework for Interrogating Binding Site Predictability and Energetic Organization of Binding Sites

To understand why certain binding sites are readily detectable while others remain elusive, we proposed a theoretical formulation that reframes the question from “how well do models predict?” to “what do prediction patterns reveal about how evolution encodes functional sites within protein energy landscapes?” The “allosteric blind spot” has typically been attributed to data sparsity or algorithmic limitations. However, an alternative possibility is that the limited detectability of allosteric sites reflects intrinsic properties of their evolutionary and energetic design. Distinguishing between these explanations requires an integrated strategy that can connect algorithmic behavior to underlying biophysical principles. To address this, we introduce an explainable AI framework that treats computational PLM predictions of binding sites not as endpoints but as diagnostic probes of the protein energy landscape organization (Figure 1). The first stream employs PLM as an unbiased evolutionary probe. When such a model encounters a binding site, its prediction confidence reports on how strongly that site is constrained by evolution-a quantitative readout of sequence-based detectability. The second stream employs the landscape-based local frustration analysis. This framework evaluates how favorably each native residue-residue interaction is optimized relative to alternative configurations, classifying residues into three energetic classes: minimally frustrated (interactions strongly favored over alternatives, reflecting evolutionary optimization), neutrally frustrated (interactions neither strongly favored nor disfavored, permitting conformational plasticity), and highly frustrated (interactions less favorable than alternatives, storing local strain).^42–47^ This provides a physically grounded language for describing how different regions of a protein are embedded within the energy landscape. The critical insight emerges from overlaying these two streams (Figure 1-figure supplement 1). If prediction performance correlates with frustration signatures, then PLM behavior becomes interpretable through protein physics rather than remaining an opaque statistical output. Specifically, we hypothesized that canonical orthosteric sites-evolutionarily locked, catalytically essential pockets-would reside in minimally frustrated basins where native interactions are strongly favored and evolutionary conservation is high, generating robust coevolutionary signals reliably detectable by PLMs. On the other hand, allosteric sites, designed for conformational plasticity and context-dependent function, would occupy shallow, rugged regions of the energy landscape dominated by neutral frustration. This neutrality permits sequence drift and structural adaptability, enabling regulatory versatility, but also erodes the statistical signals that sequence-based models require. When a PLM encounters such a site, its predictions should become more diffuse, low-confidence, and structure-dependent-a regime where the pocket is only weakly detectable in most conformations.

**Figure 1.**
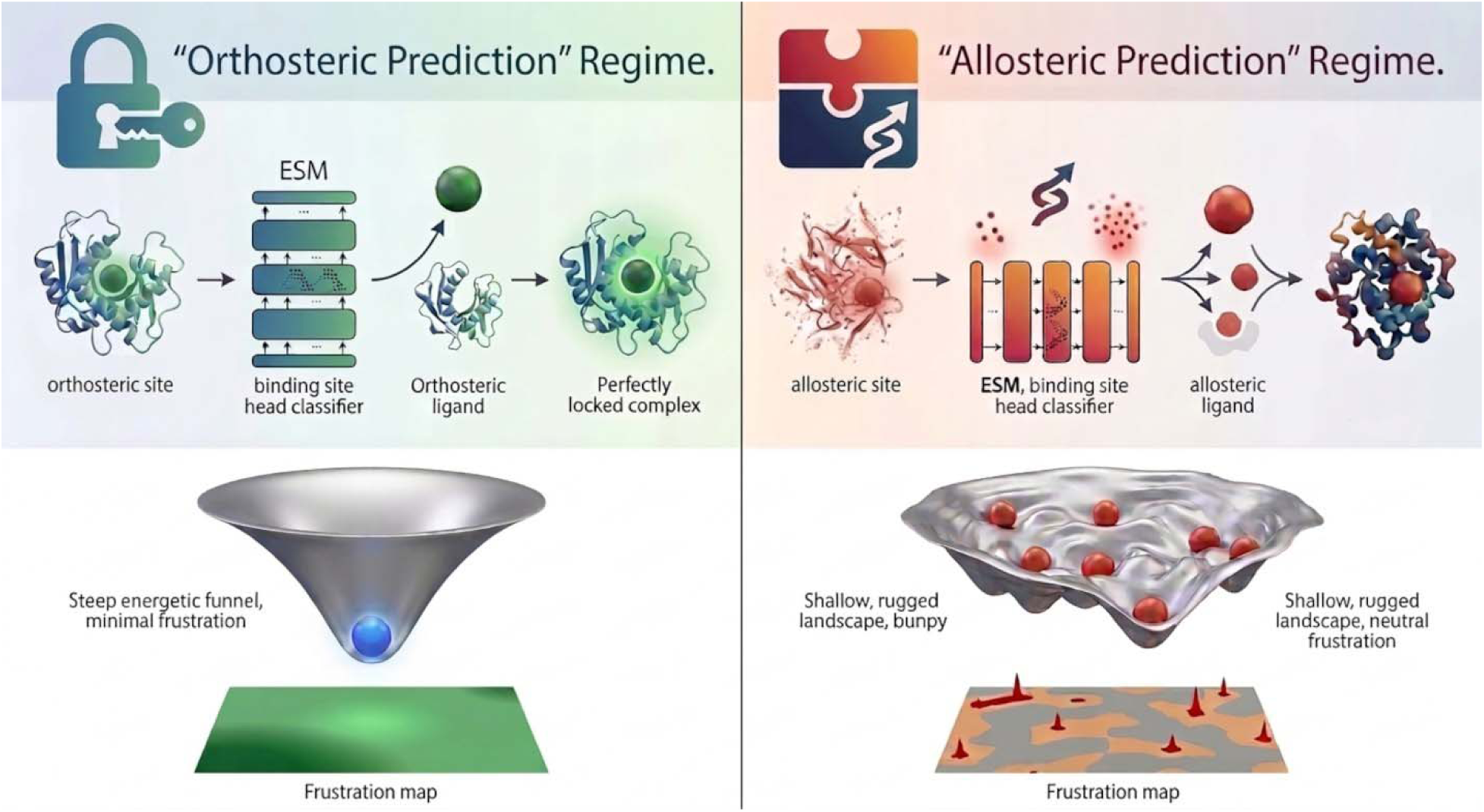
A diagnostic framework linking PLM predictions to energy landscape organization. (Left panel) Orthosteric prediction regime: PLM encounters an orthosteric pocket characterized by a steep energetic funnel with minimal frustration. The PLM detects this site with high confidence; the frustration map shows a concentrated region of minimal frustration (green) at the binding interface. (Right panel) Allosteric prediction regime: the same model encounters a dynamic allosteric pocket with a shallow, rugged landscape dominated by neutral frustration. The PLM produces diffuse, low-confidence predictions; the frustration map shows a neutrally frustrated pocket interior (red) with surrounding frustrated switch points (magenta). Bottom panels illustrate energy landscape schematics: orthosteric sites occupy deep, funneled basins that generate strong coevolutionary signals, while allosteric sites reside in shallow landscapes with multiple minima, enabling regulatory versatility at the cost of sequence-based detectability.

**Figure 1-figure supplement 1. Schematic of the unified PLM-based architecture and residue-level classification pipeline.**

Detailed representation of the computational stages used to map sequence and structural data to binding site predictions. Input and Encoding Stage (Top Panel): Protein primary sequences and 3D coordinates (from PDB or AlphaFold) are processed by a 33-layer ESM-2 transformer encoder^27^ (650M parameters, 20 attention heads). The encoder generates 1280-dimensional latent representations (hi) for every residue, capturing both local and distal evolutionary context. Classification Head Adaptation (Center Panel): The original Masked Language Model (MLM) head is replaced with a specialized linear layer (W∈R1280×1) and a sigmoid function. This architecture converts the latent residue vectors into a scalar probability (pi∈[0,1]) for binary site classification. Prediction Output and Conceptual Landscapes (Bottom Panel): Residues meeting the classification threshold (pi>0.75) are mapped to the 3D structure. The schematic energy landscapes illustrate the underlying difficulty of the task: orthosteric sites occupy deep, stable basins that yield high-confidence predictions, whereas allosteric sites-defined by rugged, multi-minima landscapes-present more challenging targets for the sequence-based model.

### Structural Diversity of Kinase Allosteric Sites Defines a Natural Testbed for Probing Allostery and Binding-Site Detectability

To investigate how evolutionary constraints shape binding site detectability, we used protein kinase family that simultaneously satisfies three criteria: a conserved catalytic architecture, extensive structural coverage across functional states, and a diverse repertoire of regulatory allosteric pockets (Figure 2).^15–19^ The KinCoRe classification system provides standardized annotation of kinase conformational states and inhibitor binding modes.^20–22^ Based on specific structural markers-the DFG motif (Asp-Phe-Gly) and the regulatory αC-helix position-this system categorizes inhibitors into five mechanistic classes (Figure 2-figure supplement 1). Type I, I.5, and II inhibitors occupy the orthosteric ATP-binding cleft but capture different conformational states: Type I binds the active DFG-in conformation, Type I.5 represents a transitional state with the αC-helix shifted outward, and Type II binds the inactive DFG-out conformation. Type III inhibitors are proximal allosteric modulators that bind pockets adjacent to the ATP site but do not enter the cleft itself, while Type IV inhibitors are distal allosteric modulators that bind sites far from the catalytic machinery (Figure 2-figure supplement 1).To visualize the diversity of allosteric sites across the kinase fold, we mapped the 12 canonical allosteric binding regions compiled from a recent survey of allosteric kinase ligands^24,25^ (Figure 2). This structural atlas reveals two important features simultaneously: significant diversity in allosteric site locations, yet recurrent patterns across the kinase family. Allosteric modulators target a wide range of locations, including the well-characterized myristoyl pocket in the C-lobe (Type IV), the back pocket adjacent to the ATP site (Type III), and several less-conserved surface grooves. This spatial diversity reflects an evolutionary strategy of exploiting multiple regulatory interfaces for fine-tuned control. Yet despite this diversity, allosteric sites consistently occupy regions peripheral to the conserved catalytic core. Only three of the binding sites of type IV inhibitors represented deep pockets (D, E, and H).^24,25^ Of special importance is deep pocket E in the C-terminal lobe that corresponds to the myristoyl pocket in ABL kinase.

**Figure 2.**
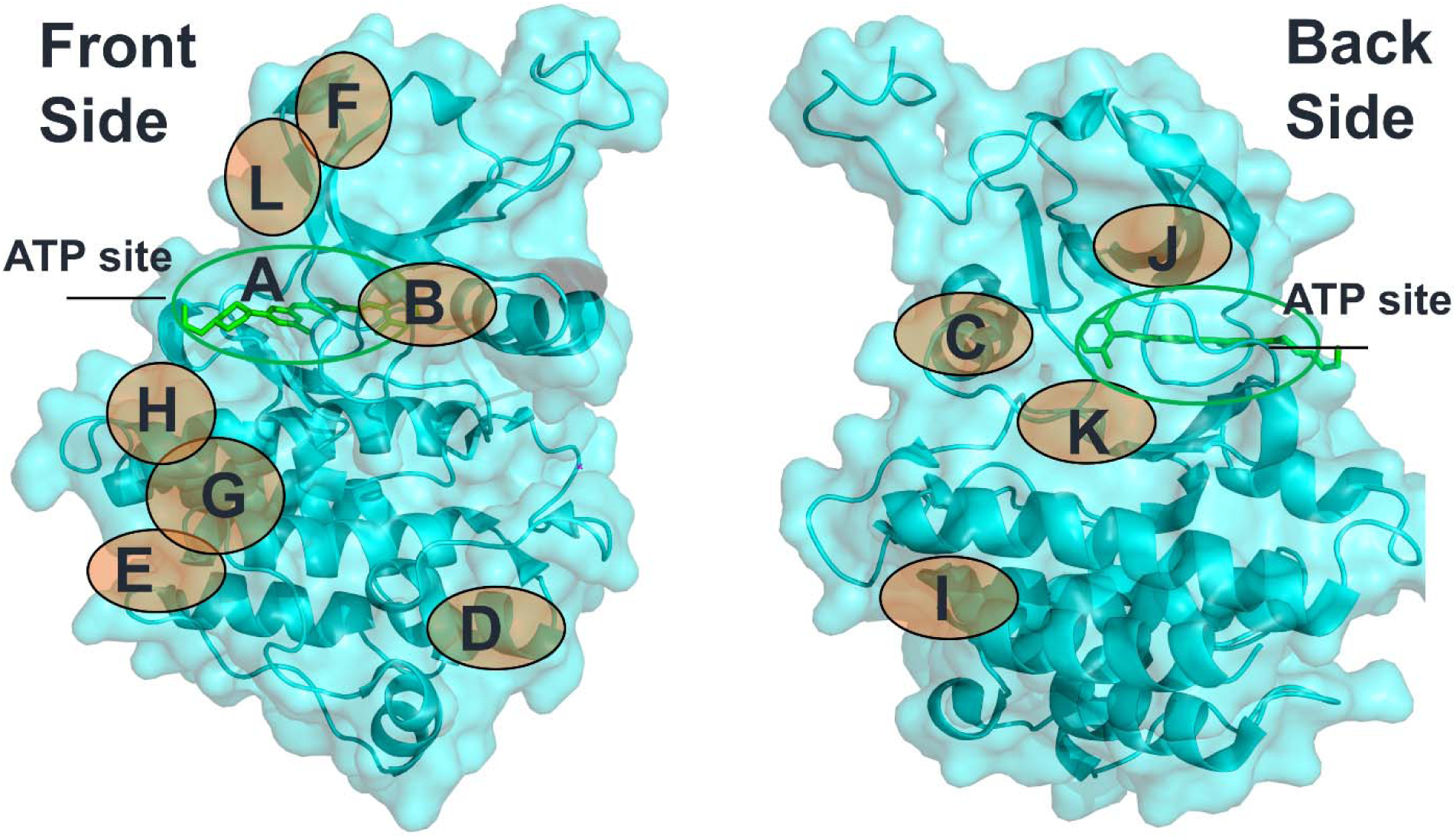
Structural diversity of allosteric binding sites across the kinase fold. The 12 canonical allosteric binding regions, compiled from a survey of 262 allosteric kinase ligands^24,25^ are mapped onto a representative ABL kinase structure (PDB 2GQG). Allosteric modulators target a wide range of locations, including the myristoyl pocket in the C-lobe (Type IV), the back pocket adjacent to the ATP site (Type III), and multiple surface grooves. The orthosteric type I Dasatinib bound in the ATP site is shown in green sticks.

Among kinase systems, ABL kinase provides a particularly informative experimental platform for probing and decoding allosteric site diversity using our explainable AI framework.^48–51^ The extensive structural characterization of ABL complexes, including a spectrum of active and inactive apo states^52,53^, regulatory domain assemblies with the SH2-SH3 module,^54–57^ and numerous ligand-bound conformations^58–63^ provides an ideal system for examining how energetic organization shapes allosteric ligand recognition. Using the ABL1 kinase as a representative architecture, we highlight the complex multi-domain organization that couples the catalytic kinase domain (KD) to regulatory SH3 and SH2 modules (Figure 2-figure supplement 2A-C). The regulatory myristoyl pocket in ABL (Figure 2) exemplifies the functional versatility of allosteric sites by functioning not as a simple regulatory switch but as a versatile allosteric hub capable of accommodating chemically diverse ligands and producing distinct functional outcomes (Figure 2-figure supplement 2). Structural studies showed that this allosteric binding pocket exists in both ligand-free apo structures and kinase complexes where some ligands can induce a significant conformational change in the pocket that triggers autoinhibition, while allosteric activators stabilize the native conformation.^24,25^ Our previous computational studies have also mapped the intricate allosteric networks in ABL kinase identifying communication pathways that connect the peripheral myristoyl binding site to the orthosteric ATP pocket (Figure 2-figure supplement 2D, E). These findings provide a structural foundation for exploring how ‘rigid’ functional motifs (minimally frustrated regions) are coupled with structurally adaptable peripheral regions (neutrally frustrated zones) - a relationship recently highlighted in broader proteomic studies (Lehner et al., 202X).. Taken together, the structural diversity of kinase regulatory pockets, combined with the conserved architecture of the catalytic site and the detailed mechanistic understanding of ABL allostery creates a natural experimental landscape and a high-resolution testbed for using PLM predictions to probe how evolutionary constraints influence binding site detectability.

**Figure 2-figure supplement 1. Structural motifs of the kinase catalytic domain and kinome-wide distribution of study targets.** (A) Representative 3D structural model of the conserved kinase fold highlighting the regulatory architecture. The Regulatory spine (R-spine) is shown in a vertically aligned, “assembled” state, a hallmark of the active conformation. The DFG motif (Asp-Phe-Gly) is positioned at the start of the activation loop (A-loop), acting as a critical conformational switch; its orientation (DFG-in vs. DFG-out) coordinates with the position of the C-helix to modulate the ATP-binding pocket’s volume and energetic accessibility. The catalytic loop provides the necessary residues for phosphotransfer. (B) Phylogenetic tree of the human kinome^64^ illustrating the breadth of the current study. Node colors categorize kinases into dark kinase domains. non-dark kinase domains and uncharacterized/non-curated domains. The model’s predictive framework was validated across all major groups-including TK, TKL, STE, CK1, AGC, CAMK, and CMGC-ensuring that the PLM-derived frustration maps and binding site classifications are robust to the evolutionary divergence of the human kinome. Kinases are obtained from the curated kinome that are visualized on the Coral kinase dendrogram.^65^ The recomputed dark kinome is shown in blue and non-dark kinases are shown in yellow. The panel was adopted from a recent study of kinome.^66^

**Figure 2-figure supplement 2. Structural organization of ABL complexes with allosteric modulators and mapping of communication pathways.** (A) Architecture of the autoinhibited ABL-SH3-SH2-KD assembly. Crystal structure of the ABL regulatory core in complex with the orthosteric inhibitor Nilotinib and the allosteric inhibitor Asciminib. The multi-domain organization is indicated: SH3 domain (dark blue), SH2 domain (green), SH2-kinase linker (cyan), and the kinase domain (KD, red). Inhibitors are shown in sticks with atom-based coloring. (B) Inactive ABL-KD conformation. Structure of the isolated kinase domain (magenta) in complex with the type II inhibitor Imatinib and the allosteric myristoyl-site inhibitor GNF-2. (C) Active-like ABL-KD conformation. Structural overlay illustrating the ABL-KD in complex with Imatinib and the allosteric activator DPH. (D–E) Spatial arrangement of allosteric communication networks identified in our previous studies^67^ that connect the allosteric myristoyl site and the orthosteric ATP-binding site. (D) Representative connectivity in the ABL1-SH3-SH2/Asciminib complex. (E) Preferential routes in the ABL1/DPH complex. Blue spheres represent the optimal pathways connecting the regulatory myristoyl site with the catalytic machinery.

### Hierarchy of Binding Site Detectability Across the Human Kinome: PLM Performance as a Diagnostic Probe of the Allosteric Blind Spot

To systematically assess binding-site detectability, we employed previously developed a fine-tuned PLM for residue-level binding site prediction.^68^ In this implementation, the pretrained ESM2-650M model^27^ was employed and optimized (Figure 1-figure supplement 1) on the human subset of the LIGYSIS dataset.^69,70^ The masked language modeling head was replaced with a token-level classification layer that maps each residue’s 1280-dimensional latent representation to a calibrated probability of ligand involvement (Figure 1-figure supplement 1). Using this fine-tuned PLM in the current study as an unbiased algorithmic probe, we evaluated its predictions of binding sites across all 453 human kinases in the KinCoRe dataset^21^ which provides rigorous geometry-based classification of inhibitor-bound structures into five mechanistic classes: Type I, I.5, and II (orthosteric ATP-competitive inhibitors) and Type III and IV (proximal and distal allosteric modulators).^68^ For each structure, we computed standard metrics AUROC, AUPR, MCC, accuracy, and F1 score-to quantify the PLM ability to identify ligand-binding residues. The resulting distributions are shown in Figure 3.

**Figure 3.**
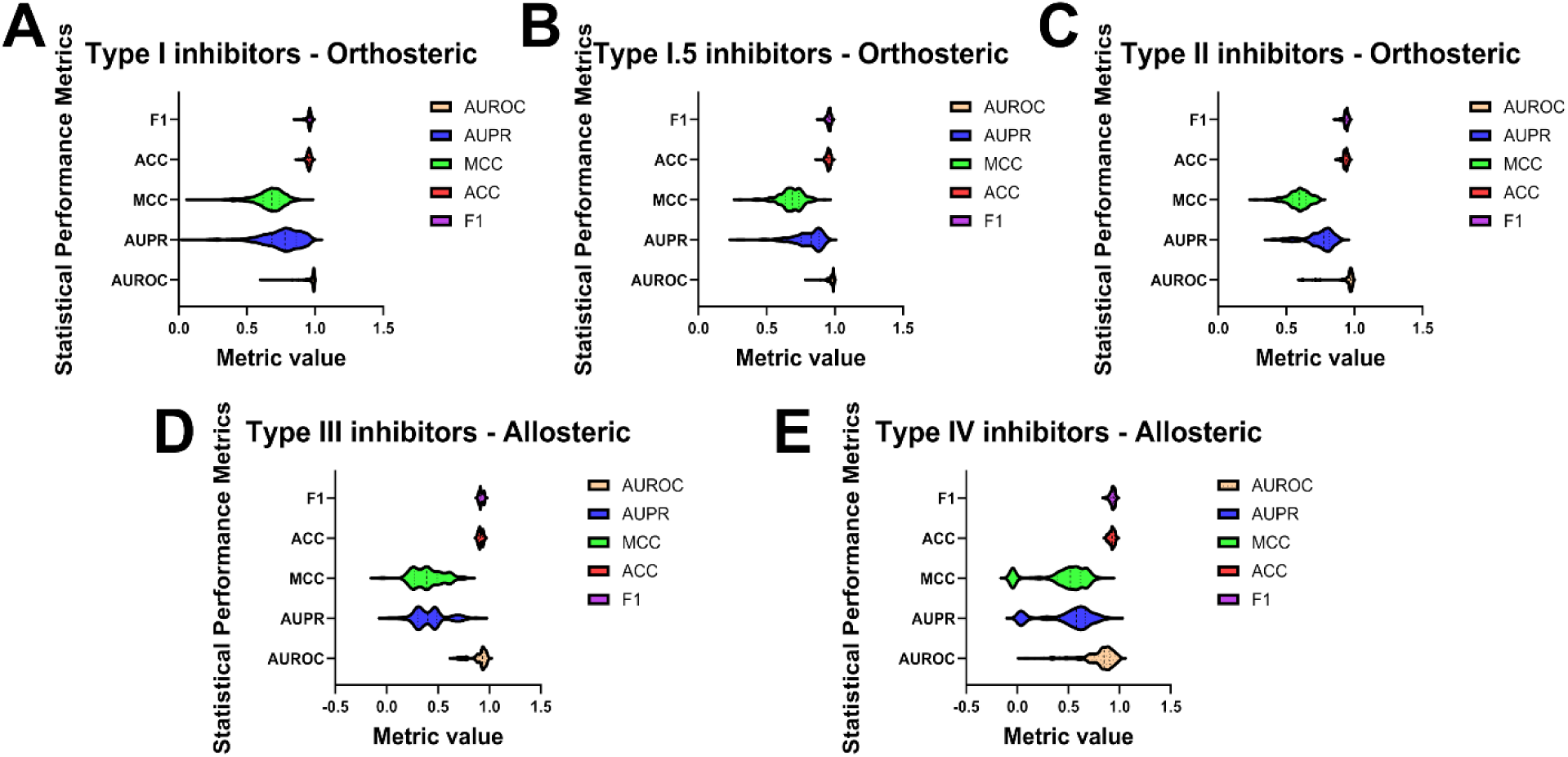
PLM analysis across orthosteric and allosteric kinase sites reveals a systematic predictability gap . Violin plots display the distribution of five classification metrics (AUROC, AUPR, MCC, Accuracy, and F1 score) for PLM predictions across five inhibitor classes from the KinCoRe dataset. Each violin represents the density of per-structure performance values, with overlaid box plots indicating median and interquartile range. The violin plots illustrate the distribution of performance metrics across structures. (A–C) Orthosteric ATP-competitive inhibitors: Type I (A), Type I.5 (B), and Type II (C) inhibitors exhibit tight distributions with median values near 1.0 and minimal interquartile ranges (IQR ≈ 0.02–0.03), indicating universal detectability across all kinase structures. (D–E) Allosteric inhibitors: Type III (D) and Type IV (E) allosteric modulators show dramatically broader, lower-performance distributions. Type III sites display moderate-to-broad distributions (median ≈ 0.36 for AUPR), reflecting structure-dependent variability, while Type IV sites exhibit extreme distribution spread (median AUPR ≈ 0.08), indicating statistical invisibility in most conformations. As the PLM outputs for each residue a probability *p*_i_ ∈ [0,1] that the residue belongs to a ligand-proximal site, we defined threshold the probabilities to obtain binary predictions. In the *PLM fixed* setting, the probabilities were binarized using a single threshold *t* = 0.75. Raw probabilities were retained for threshold-free analyses using AUROC and AUPR. The classification metrics (MCC, Accuracy, F1) were calculated using a consistent fixed confidence threshold to ensure uniformity across diverse structural contexts.

Noteworthy, while our initial methodological study^68^ introduced the fine-tuned PLM and benchmarked the predictive PLM performance and algorithmic parameters, here we deploy these metrics as diagnostic signatures and quantitative readouts revealing how the protein energy landscape can dictate the encoding of orthosteric sites versus allosteric binding pockets. The AUROC metric, which captures global ranking ability independent of decision thresholds, immediately distinguishes the three classes. Orthosteric Type I, I.5, and II sites exhibit remarkably narrow and tall distributions, with medians between 0.94 and 0.98 and interquartile ranges of only 0.02–0.03. This means that almost every individual structure was predicted well (high density at high scores), reflected in tight distributions with consistent median. This exceptional consistency indicates that orthosteric sites are universally rankable across all kinase structures-the model distinguishes them from background regardless of conformational state (Figure 3A-C).

For proximal Type III allosteric sites, the AUROC distribution broadens considerably (median ≈ 0.91, IQR ≈ 0.06–0.08) (Figure 3D,E). This widening reflects a pronounced structure-dependent variability whereby some protein kinase conformations allow near-orthosteric ranking, while the majority of predictions yield substantially lower values. The distribution is no longer a tall peak but a wider mountain, signaling that detectability is contingent on structural context rather than universal. Distal Type IV allosteric sites display the most extreme pattern: an extremely broad, flattened distribution (median ≈ 0.68, IQR ≈ 0.12–0.15) with many structures yielding near-random ranking and a long lower tail extending to 0.5 (Figure 3E). This spread reveals that Type IV sites are statistically invisible in most conformations, with detectable signal present only in a subset of structures.

**Figure 3-source data 1. Distribution statistics of performance metrics across kinase site types.** This Excel workbook contains statistical summaries for model performance, including AUROC (Table S1), AUPR (Table S2), MCC (Table S3), and weighted F1 scores (Table S4). A summary of performance metric characteristics and their sensitivity to class imbalance is also provided (Table S5).

The AUPR metric, which measures operational precision under extreme class imbalance, exposes the true extent of the “allosteric blind spot.” Orthosteric sites maintain usable precision across most structures, with moderately broad but right-skewed distributions (median AUPR 0.63–0.75, IQR ≈ 0.08–0.10) (Figure 3A-C). The skew reflects class imbalance rather than site ambiguity-some structures yield slightly lower recall, but precision remains robust. Type III sites show broader AUPR distributions (median ≈ 0.36, IQR ≈ 0.15–0.20), spanning from approximately 0.10 to 0.60 (Figure 3D). This extreme width indicates that precision is highly structure-dependent where certain conformations “reveal” the site with moderate precision, while most perform poorly. For Type IV sites AUPR distributions are compressed near zero, with medians of only 0.08 and spreads that place most structures near the random baseline of 0.02–0.08 (Figure 3E). The characteristic pattern of high AUROC coupled with vanishing AUPR for Type IV sites reveals that while allosteric residues may be globally rankable, they lack the discriminative evolutionary signatures required for confident detection, reflecting their design for conformational plasticity rather than conserved recognition.

MCC, the most informative metric for imbalanced data when using a fixed confidence threshold, confirms and refines this hierarchy. Orthosteric sites demonstrate tight, high distributions (median MCC 0.58–0.69, IQR ≈ 0.05–0.07), with a sharp peak in the 0.6–0.7 range and very few structures falling below 0.4 (Figure 3A-C). This consistency reflects high-quality classification across kinase structures. Type III sites exhibit broader, lower distributions (median MCC 0.38–0.45, IQR ≈ 0.10–0.12) (Figure 3D). While some structures still achieve MCC above 0.5, many fall to 0.2–0.3, with the distribution spanning from near-zero to moderately positive-a direct reflection of conformational ambiguity. Type IV sites collapse toward zero (median MCC < 0.15, IQR ≈ 0.12–0.15), with most mass lying between –0.1 and +0.2 and only a long tail extending to perhaps 0.3–0.4 for rare outliers (Figure 3E). Many structures yield MCC values indistinguishable from random, confirming statistical invisibility. Accuracy, which is influenced by the large number of true negatives, shows only subtle differentiation: orthosteric sites achieve very high median accuracy (0.94–0.96) with extremely tight distributions (IQR ≈ 0.01–0.02) (Figure 3A-C); Type III sites show a slight downward shift and increased spread (median 0.88–0.92, IQR ≈ 0.03–0.04) (Figure 3D); and Type IV sites exhibit a longer lower tail (median 0.86–0.90, IQR ≈ 0.04–0.06), indicating that false positives become numerous enough to degrade accuracy in some structures (Figure 3E). Summary of distribution statistics for kinome-wide performance metrics are provided in Figure 3—source data 1. Our model demonstrated high ranking performance across orthosteric sites, while showing a significant performance decay in Type IV allosteric regions (Figure 3-source data 1). While AUROC remains relatively high for allosteric sites, the precipitous drop in AUPR and MCC confirms that these sites are statistically ‘blind’ in most Type IV allosteric sites as shown in Tables S2, S3 (Figure 3-source data 1). While accuracy alone does not reveal the allosteric blind spot as it is dominated by true negatives, the pattern remains consistent. The weighted F1 score, sensitive to positive-class performance, captures the precision collapse more vividly: orthosteric sites maintain high F1 across structures (median 0.94–0.96, IQR ≈ 0.03–0.05); Type III sites show very broad distributions (median 0.86–0.91, IQR ≈ 0.08–0.10); and Type IV sites, despite modestly high medians (0.85–0.89), exhibit a lower tail extending below 0.7 for many structures (Figure 3). This is the operational manifestation of the allosteric blind spot-false positives overwhelm true positives, dragging down the very metric designed to balance them.

The shapes of these distributions revealed reproducible signatures that differentiate site classes. The tight, tall distributions of orthosteric sites indicate that these sites are universally detectable-their signal is strong and invariant across structures. The broader, more variable distributions of Type III sites suggest that detectability depends on structural context. The extremely broad, collapsed distributions of Type IV sites reveal that these sites are only weakly detectable in most structures. These observations raise a fundamental question: What physical property of the sites themselves could explain these systematic differences in algorithmic detectability? The patterns we observe-tight versus broad versus collapsed distributions-suggest differences in how consistently these sites are encoded in sequence information. Orthosteric sites appear to be strongly and invariantly encoded while allosteric sites appear to be weakly and variably encoded. This finding points toward the need for a biophysical interpretability layer-a framework that can connect algorithmic behavior to underlying physical properties of protein structures. In the following sections, we introduce local frustration analysis as such a framework and test directly whether the performance patterns observed here correspond to differences in how these sites are energetically embedded within the protein energy landscape.

### Local Frustration Analysis Reveals Landscape-Encoded Determinants of Prediction Dichotomy Between Orthosteric and Allosteric Binding Sites

We conjectured that the limitations of the PLM in detecting allosteric binding sites arise from a fundamental consequence of the intrinsic biophysical design of these sites. To address this and mechanistically explain the divergent performance of the PLM on orthosteric versus allosteric kinase binding sites, we introduce local frustration analysis as a biophysical interpretability layer-a framework that connects algorithmic behavior to underlying physical properties of protein structures. Frustration analysis partitions residue–residue interactions into three energetic classes-minimally frustrated, neutrally frustrated, and highly frustrated based on how the native interaction energy compares to an ensemble of decoys generated either by randomizing residue identities (mutational frustration) or by perturbing both identities and local geometry (conformational frustration).^42–47^ For minimally frustrated contacts, native interactions strongly favored over alternatives; neutrally frustrated are characterized by native interactions that are neither strongly favored nor disfavored; and for highly frustrated contacts, native interactions are less favorable than alternative decoys pointing to significant local strain.^42,43^

Here we test the hypothesis that predictive success is governed by the underlying frustration signature of a functional site. We began by examining the global frustration landscape of the kinase domain. For each structure in our dataset, we computed both conformational frustration (sensitivity to structural perturbations) and mutational frustration (sensitivity to amino acid substitutions) and visualized the distributions across all residues (Figure 4A-D). Strikingly, the global frustration profiles of orthosteric and allosteric kinase complexes were nearly indistinguishable in both conformational frustration (Figure 4A,C) and mutational frustration regimes (Figure 4B,D). Conformational frustration distributions show a dominant, broad peak centered in the neutral frustration regime, indicating that most interactions throughout the kinase fold are neither strongly optimized nor strongly strained. The violin plots are wide and roughly symmetric, reflecting the heterogeneous mix of structural elements that comprise the kinase domain (Figure 4A,C). A similar pattern emerges for mutational frustration where the majority of residues also fall within the neutral regime, with the distributions tapering off gradually toward both minimal and highly frustrated extremes (Figure 4B,D). This similarity reveals that the global energetic architecture of the kinase catalytic domain is largely conserved. The kinase domain fold provides a shared platform where neutral frustration predominates, enabling the conformational flexibility required for transitions between active and inactive states.

**Figure 4.**
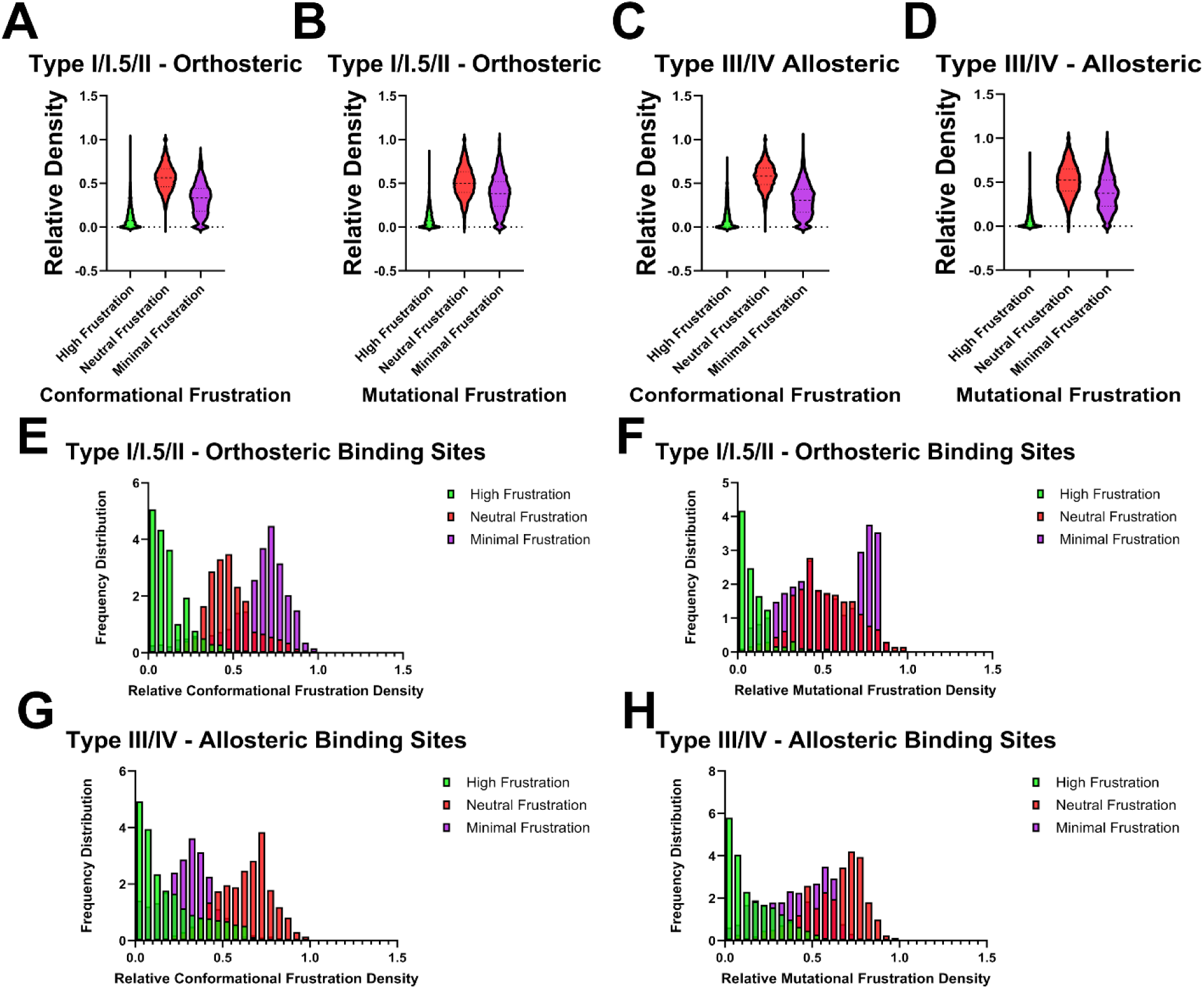
Distinct frustration landscapes encode the predictability of orthosteric versus allosteric kinase binding sites. (A–D) Global frustration architecture of the kinase domain. Violin plots show the relative density of frustration classes for the entire kinase domain fold, independent of binding site location. Panels display conformational frustration (A, C) and mutational frustration (B, D) for complexes with orthosteric inhibitors (A, B) and allosteric modulators (C, D). The distributions are nearly superimposable across both classes, dominated by neutral frustration (red), indicating that the global energetic scaffold of the kinase is conserved and intrinsically plastic regardless of ligand type. (E–H) Binding site-specific frustration distributions are show in color-coded filled bars : green for high frustration (energetically strained); red for neutral frustration (energetically indifferent); and purple for minimal frustration (energetically optimized). Histograms display the frequency distribution of relative frustration density for residues strictly within annotated ligand-binding pockets. Frustration profiles for orthosteric binding sites (E, F) exhibit a pronounced skew toward minimal frustration (purple), particularly for mutational frustration (F), where the distribution peaks at high densities (>0.7). In stark contrast, allosteric binding sites (G, H) are overwhelmingly dominated by neutral frustration (red) with a minor but consistent population of high frustration (green). The near-absence of minimal frustration in allosteric pockets indicates evolutionary permissiveness and conformational adaptability, explaining their statistical invisibility to sequence-based models.

When we restrict analysis to residues within annotated binding pockets, a strikingly different picture emerges, and here the shapes of the distributions become powerfully diagnostic. Orthosteric binding sites exhibit distributions that are sharply shifted toward minimal frustration, with a pronounced peak at high minimal frustration densities (>0.7) (Figure 4E,F). The histogram peaks sharply in the minimal frustration regime, with relative densities reaching their maximum at high minimal frustration values (>0.7) and tapering off rapidly in the neutral and highly frustrated regimes (Figure 4E). This is the signature of intense purifying selection: evolution has locked these residues in place because catalytic function demands precise geometry. The enrichment is even more dramatic for mutational frustration (Figure 4F), where the distribution is almost entirely confined to the minimal regime, reflecting the strong destabilization that would result from amino acid substitutions at these positions.

**Figure 4-figure supplement 1. Structural mapping of high neutral frustration density across diverse kinase-allosteric inhibitor complexes.** To examine the relationship between evolutionary constraints and allosteric sites, we mapped the regions of high neutral frustration density >0.7 (shown in blue) onto diverse kinase structures with type III and type IV allosteric inhibitors. Protein molecular surfaces are colored by structural origin: pale pink (Chain A) or cyan (Chain B). The corresponding allosteric inhibitors (sphere representation, atom-based coloring) are shown bound to different regulatory pockets, encompassing both Type III (adjacent to the ATP site) and Type IV (distal) modes of allosteric modulation. (Top Row) Type III adjacent-pocket inhibition. High neutral frustration density is seen localized around the orthosteric C-helix/DFG region and adjacent pockets for c-SRC (pdb 3F3U), ABL (pdb 3QRK), IGF1R (3LW0) and FAK kinase (pdb 4I4F) (Bottom Row) Distal and Type IV allosteric inhibitors. Neutral frustration density in complexes with Type IV allosteric inhibitors CDK2 (3PXF), CHK1 (pdb 3JVS), PDK1 (4AW1) and JNK1 (pdb 3O2M). These neutrally frustrated patches illustrate the structurally diverse “allosteric blind spots.”

Conformational frustration regime for allosteric binding sites (Figure 4G) exhibits a fundamentally different histogram shape. Here the distribution peaks squarely in the neutral frustration regime (0.65–0.75), with minimal frustration densities markedly reduced compared to orthosteric sites. The histogram is broader and more evenly distributed, indicating that allosteric sites are evolutionarily permissive-many substitutions are energetically tolerated, enabling sequence drift and conformational plasticity without compromising function. Mutational frustration distributions for allosteric site residues (Figure 4H) reinforce and amplify this dichotomy.

While for orthosteric sites the mutational frustration histogram is almost entirely confined to the minimal regime, the distribution for allosteric sites shifts dramatically toward neutral frustration, with minimal frustration nearly absent. Notably, both conformational and mutational frustration histograms for allosteric sites show small but consistent tails extending into the highly frustrated regime. The contrasting shapes of the frustration histograms for orthosteric and allosteric binding site residues suggests a plausible physical rationale for the observed PLM performance patterns. The sharp peaks of orthosteric sites correspond to deep, funneled energetic basins that generate strong, unambiguous coevolutionary signals PLMs detect with high confidence. The broad, neutral-dominated histograms of allosteric sites reflect shallow, rugged landscapes where evolutionary signals are likely weak and variable, precisely matching the diffuse, low-confidence and structure-dependent detectability observed for Type III and IV sites.

### Architectural Patterns of Neutral Frustration Across the Kinome

The quantitative dichotomy raises two interrelated questions about spatial organization: Where are allosteric sites located across the kinase fold? and Where does neutral frustration localize within kinase structures? To address these questions, we performed complementary mapping analyses that together reveal a striking convergence. To visualize the structural distribution of neutral frustration, we mapped high-density neutrally frustrated positions onto the kinase domains of multiple representative kinases bound to various allosteric ligands (Figure 4-figure supplement 1). Each panel displays a different kinase–ligand complex, with neutrally frustrated residues rendered in blue. Despite the diversity of kinases and ligands, a consistent architectural pattern emerges that directly parallels the allosteric site map (Figure 4-figure supplement 1). The ATP-binding cleft, the canonical activation segment, and the central hydrophobic spine residues essential for catalysis exhibit minimal frustration density, consistent with their evolutionary constraint: these elements must maintain defined conformations to support phosphotransfer activity and are therefore under strong purifying selection. Neutral frustration consistently localizes to peripheral and conformationally adaptable regulatory regions-flexible loops, dynamic hinges, and lineage-specific pockets-while being systematically excluded from the conserved catalytic core (Figure 4-figure supplement 1). Thie extended mapping of high neutral frustration density on representative kinase complexes with type III and type IV allosteric inhibitors reinforces the notion that regulatory potential may be encoded in regions of the energy landscape that are evolutionarily permissive and conformationally plastic.

Across all complexes, neutral frustration consistently decorates three classes of regions: flexible regulatory loops (activation loop, P+1 motif), dynamic mechanical hinges (αC-helix, DFG motif, lobe connector), and lineage-specific allosteric pockets (myristoyl site, DFG-out pocket, C-lobe regulatory interfaces) (Figure 4-figure supplement 1). This corresponds to the location of several characterized allosteric binding interfaces, and the neutral frustration of the P+1 motif likely reflects its need to accommodate diverse substrate sequences while maintaining recognition specificity. Similarly, neutral frustration consistently decorates the hinges that connect rigid elements, such as the αC-helix which swings between “in” and “out” conformations to control catalysis, and the DFG motif which flips to gate ATP and inhibitor binding (Figure 4-figure supplement 1). The key finding of this analysis is that the neutral frustration density overlaps significantly with both proximal (Type III) and distal (Type IV) allosteric sites. In ABL the myristoyl pocket in the C-lobe is clearly delineated by high neutral frustration density (Figure 4-figure supplement 1). A significant neutral frustration in the allosteric pockets reflects this evolutionary history: they remain plastic, enabling each kinase family to develop specialized regulatory interfaces while preserving the underlying conformational flexibility required for function. The neutral frustration density is not necessarily connected as a single continuous surface. In many structures, it appears as multiple distinct small clusters distributed across the kinase domain-a cluster in the A-loop, another around he αC-helix hinge, a third in the myristoyl pocket. The minimally frustrated catalytic core must remain rigid to maintain catalytic geometry, while neutral frustration is strategically deployed at regulatory junctions where conformational transitions occur. This creates an energetic mosaic: stable cores interspersed with flexible switches. This discontinuous organization suggests that neutral frustration is a local feature of specific structural elements that have evolved for conformational flexibility. Yet these spatially separated neutral frustration clusters do not function in isolation; rather, they form interconnected nodes within allosteric communication networks that couple distal regulatory sites to the catalytic center.

Drawing from our previous studies of allosteric interaction networks in protein kinase structures and complexes^71–76^ as well as other theoretical studies^77^ and a significant body of NMR experiments^78–80^ the discontinuous but organized distribution of frustration clusters can reflect their integration into allosteric communication networks: stable positions provide reliable signal transmission pathways, while neutrally frustrated residues enable the conformational flexibility required for network switching. This hybrid architecture-rigid backbone elements interspersed with frustrated switching points-transforms local ligand binding into global regulatory outcomes, allowing the kinase to integrate diverse allosteric signals into a coherent functional response.

While neutral frustration dominates the stable kinase conformations, our previous studies also revealed a dynamic dimension to this architectural principle. We previously found that localized highly frustrated clusters in inactive kinase states could collocate with the regions directly involved in conformational changes associated with kinase function.^71,72^ This suggests a dual role for frustration in these regulatory regions. Neutral frustration establishes the baseline thermodynamic landscape that permits conformational sampling within stable functional states-this is the energetic signature of the “permissive scaffold” we observe across the kinase domain. However, localized clusters of high frustration within these neutrally frustrated regions serve as conformational nucleation points that become particularly pronounced in higher-energy inactive states, creating the strain necessary to drive large-scale structural transitions. Taken together, these findings suggest that allosteric sites are encoded in neutrally frustrated regions optimized for context-dependent regulatory modulation while the capacity to accumulate high frustration in transition states provides an energetic driving force for conformational switching. This duality explains both the remarkable regulatory versatility of kinase allosteric sites and their systematic invisibility to current predictive algorithms.

### ABL Kinase as a Canonical System for Decoding Frustration Signatures

The previous sections established two complementary findings at the kinome-wide level. First, orthosteric and allosteric sites exhibit systematically different patterns of PLM detectability (Figure 3). Second, these differences in algorithmic behavior reflect underlying differences in energetic embedding: orthosteric sites are enriched in minimal frustration, while allosteric sites are dominated by neutral frustration and localize to peripheral regions of the kinase fold (Figure 4).

To anchor the frustration trends in a physically interpretable molecular system, we now examine these principles at atomic resolution using the ABL kinase system where the relationship between neutral frustration, conformational plasticity, and regulatory versatility can be explored in molecular detail across multiple ligand-bound states and regulatory contexts. The regulatory myristoyl pocket in ABL (Figure 2) exemplifies the functional versatility of allosteric sites: the clinically approved drug Asciminib binds here to stabilize an autoinhibited conformation^59,60^; GNF-2 and GNF-5 also target this pocket but function as inhibitors^61^; the small molecule DPH acts as an activator^62^; and the ATP-competitive inhibitor imatinib itself can bind to this pocket, where the interaction both competes with catalytic inhibition and promotes conformations associated with increased kinase activity.^63^ The depth of experimental characterization enables us to test whether the principles inferred from kinome-wide analysis hold up under detailed atomic scrutiny.

Residue-level frustration profiling was performed across an ensemble of ABL complexes spanning active, intermediate, and fully autoinhibited states bound to orthosteric and allosteric ligands, providing a structural-scale test of the hypothesis that orthosteric and allosteric sites are encoded in fundamentally different energetic regimes (Figure 5). In ABL complexes bound to type I ATP-competitive inhibitors, the orthosteric binding cleft is strongly enriched in minimally frustrated residues, both conformationally and mutationally (Figure 5A). The probability density is sharply shifted toward the minimal-frustration regime, with a pronounced population at high minimal-frustration fractions (0.7–0.9), while maintaining a modest neutral-frustration background. This pattern defines an energetically locked interaction hotspot in which native residue identities and geometries are highly optimized and substitutions are strongly penalized.

**Figure 5.**
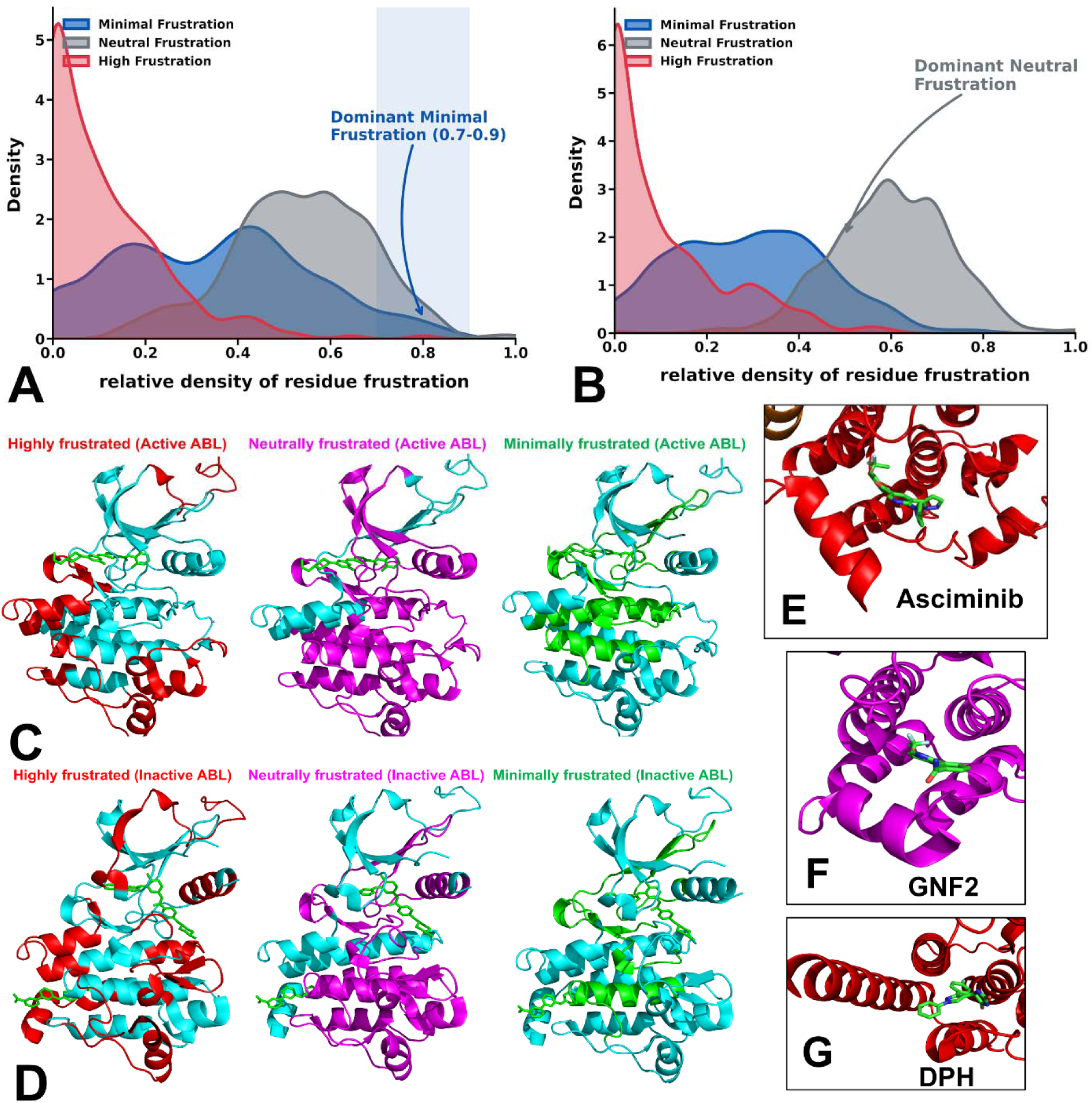
Atomic-resolution frustration signatures of orthosteric and allosteric sites in ABL kinase reveal distinct energetic and structural fingerprints. (A) Cumulative probability density of conformational frustration for orthosteric binding site residues aggregated over all ABL complexes with orthosteric inhibitors. A pronounced high-density region between 0.7 and 0.9 on the x-axis (blue-shaded background) is dominated by minimally frustrated interactions (deep blue curve), indicating strong energetic optimization of the ATP-binding cleft. (B) Cumulative probability density of conformational frustration for allosteric binding site residues aggregated over all ABL complexes with allosteric inhibitors. The distribution is dominated by neutrally frustrated interactions (grey curve) and exhibits a shift toward higher frustration values, reflecting energetic permissiveness and conformational heterogeneity. (C) Structural mapping of local frustration hotspots in the ABL complexes bound to orthosteric type I/II inhibitors (ligand shown in sticks; kinase domain in light pink ribbons). Highly frustrated hotspots (top 10%, red ribbons) localize predominantly to flexible regulatory regions, including the activation loop. Neutrally frustrated hotspots (top 10%, magenta ribbons) form a percolating scaffold throughout the fold, supporting conformational adaptability. Minimally frustrated hotspots (top 10%, green ribbons) cluster in the conserved ATP-binding cleft and catalytic core, coinciding directly with the orthosteric binding site. (D) Structural mapping of frustration hotspots in inactive ABL complexes bound to the type II orthosteric inhibitor and ABL allosteric inhibitors (both ligands shown in sticks). Highly frustrated hotspots (red ribbons) concentrate in regulatory switch regions. Neutrally frustrated hotspots (magenta ribbons) permeate the kinase domain and are enriched around the myristoyl allosteric pocket. Minimally frustrated hotspots (green ribbons) localize in the catalytic core and orthosteric site. (E) Close-up view of the myristoyl allosteric pocket bound by the allosteric inhibitor Asciminib (ligand in atom-colored sticks) showing pronounced bending of the αI-helix associated with stabilization of the ABL autoinhibited conformation. (F) Close-up view of the allosteric pocket bound by the inhibitor GNF-2, highlighting a distinct ligand-induced deformation that stabilizes an inactive conformation. (G) Close-up view of the myristoyl pocket bound by the allosteric activator DPH, showing partial unbending of the αI-helix that disrupts autoinhibitory constraints and promotes active ABL conformation.

This architecture reflects strong evolutionary constraint and geometric recurrence, producing the conserved sequence and structural signatures that PLMs and structure-based predictors recognize with high precision. In contrast, allosteric sites in ABL in complexes with allosteric modulators exhibit a fundamentally different frustration signature (Figure 5B). The probability density shifts toward neutral frustration, with a broader distribution and an increased contribution from highly frustrated residues. The results reflect spatial distribution of allosteric pockets embedded within a scaffold of predominantly neutrally frustrated residues, punctuated by localized clusters of high frustration at regulatory junctions involving the αI-helix, activation loop, and regulatory spine (Figure 5B).

**Figure 5-figure supplements 1–5: General Legend.**

For all frustration profiles, data are categorized by frustration index: Minimally Frustrated (MF, blue diamonds), Neutrally Frustrated (NF, red squares), and Highly Frustrated (HF, grey bars). Profiles are generated using Configurational (spatial) and Mutational (evolutionary) decoy sets.

**Figure 5-figure supplement 1. ABL kinase with Dasatinib (PDB: 2GQG).**

Residue-level frustration for the kinase domain (residues 220–510) bound to a Type I inhibitor. (A) Configurational Frustration: The ATP-binding pocket and hydrophobic core exhibit high MF density, indicating structural stabilization. NF and HF residues are restricted to peripheral loops. (B) Mutational Frustration: Strong agreement between MF clusters confirms the orthosteric site is both energetically and evolutionarily optimized, establishing a “locked” baseline.

**Figure 5-figure supplement 2. ABL kinase with Axitinib (PDB: 4AW9).**

Validation of orthosteric signatures using the Type I inhibitor Axitinib (residues 230–510). (A) Configurational Frustration: Dense MF interactions within the catalytic machinery reflect a stabilized active-like conformation. Gaps (residues 275–295) denote crystallographic disorder. (B) Mutational Frustration: Preservation of MF clusters confirms that the orthosteric pocket is evolutionarily optimized, providing a reference for comparison with allosteric regions.

**Figure 5-figure supplement 3. Autoinhibited ABL-SH3-SH2-KD complex (PDB: 5MO4).** Analysis of the full regulatory assembly bound to Nilotinib (orthosteric) and Asciminib (allosteric). (A) Configurational Frustration: Widespread MF clusters define the domain interfaces of this “clamped” state (residues 80–540). Notably, the myristoyl pocket remains dominated by NF residues despite Asciminib occupancy. (B) Mutational Frustration: Persistent NF density confirms that the allosteric site maintains a neutral energetic signature independent of ligand-induced stabilization.

**Figure 5-figure supplement 4. ABL kinase with Imatinib and GNF-2 (PDB: 3K5V).** Analysis of the kinase domain (residues 230–530) with dual occupancy of the ATP (Imatinib) and myristoyl (GNF-2) pockets. (A) Configurational Frustration: The dual inhibitors stabilize the inactive DFG-out conformation. While the core shows significant MF clusters, the allosteric myristoyl pocket and surrounding network maintain high NF density. (B) Mutational Frustration: Persistent NF density confirms the lack of rigid evolutionary optimization at the allosteric interface.

**Figure 5-figure supplement 5. ABL kinase with Imatinib and the activator DPH (PDB: 3PYY).**

Analysis of the kinase domain (residues 240–530) bound to an allosteric activator. (A) Configurational Frustration: Unlike allosteric inhibitors, the activator DPH results in a heterogeneous landscape. While the orthosteric pocket remains stable (MF), the allosteric site exhibits high NF and HF density, reflecting the energetic strain required to disrupt autoinhibition. (B) Mutational Frustration: The allosteric machinery maintains high NF density even under activation, highlighting the shared neutral signature of regulatory pockets.

Structural mapping of frustration hotspots in the ABL orthosteric complexes reinforces this interpretation (Figure 5C). Minimally frustrated residues concentrate in the catalytic core, including the central β-sheet, DFG motif, HRD catalytic loop, and regulatory spine, defining a stability nucleus that enforces a conserved geometry for ATP binding and catalysis. Neutrally frustrated residues form a continuous scaffold linking the N- and C-lobes, providing flexibility for hinge motion and αC-helix repositioning. Highly frustrated residues cluster at known conformational switch regions, including the activation loop and αC-helix interface, where strain is locally stored to facilitate state transitions. Thus, orthosteric recognition is embedded in a landscape that is globally plastic but locally locked at the binding site. Structural projection of frustration classes in the ABL allosteric complexes (Figure 5D) shows that highly frustrated residues form a contiguous frustrated region involving the A-loop (residues 384–410), which folds inward to make extensive contacts with the αG-helix (residues 425–445), and the intervening P+1 motif (residues 404–412). This tripartite assembly is characterized by non-native residue-residue contacts and energetic strain, and it plays a critical role in allosteric control by locking ABL in the inactive state. These highly frustrated clusters correspond to regions that undergo large structural rearrangements during inactive-to-active transitions, effectively serving as “initiation cracking points” that can perturb the inactive state and promote functional switching.^81,82^ This analysis reinforces the notion of a dynamic frustration model in which dynamic regulatory regions that exhibit neutral frustration in dormant, low-energy states could accumulate high frustration in more strained inactive conformations. We argue that this conformational dependence of frustration signatures has important implications for understanding allosteric site detectability. Allosteric pockets reside in regions that sample multiple frustration regimes across the conformational ensemble. In some states, these regions display neutral frustration (permitting conformational sampling); in others, they accumulate high frustration (priming transition). This frustration heterogeneity across states further erodes the consistent evolutionary signals that PLMs require for reliable detection. By contrast, orthosteric sites maintain minimal frustration across all conformational states, reflecting their role as stable anchoring points that must preserve catalytic geometry regardless of regulatory state. This frustration stability generates the robust coevolutionary signals that enable high-precision detection.

To provide a more granular, atomic-resolution illustration of the general frustration signatures described for orthosteric and allosteric binding sites, we present a detailed residue-level analysis of high, neutral, and minimally frustrated positions across distinct ABL kinase complexes : (a) the active and inactive catalytic domain conformations formed by type I kinase inhibitors Dasatinib (pdb id 2GQG)^83^ and Axitinib (pdb id 4WA9)^84^ ; (b) the inactive ABL structure bound with type II inhibitor Imatinib and allosteric inhibitor GNF-2 (pdb id 3K5V)^61^, inactive ABL complex with type II Imatinib and allosteric activator DPH (pdb id 3PYY)^62^ and the autoinhibitory ABL regulatory complex with type II inhibitor Nilotinib and allosteric inhibitor Asciminib (pdb id 5MO4)^59,50^. The residue-based frustration profiling of ABL kinase complexes with the type I inhibitors Dasatinib (Figure 5-figure supplement 1) and Axitinib (Figure 5-figure supplement 2) provides a specific, atomic-resolution illustration of the general frustration signatures described for orthosteric binding sites. Conformational frustration and mutational frustration profiles for these complexes display the canonical energetic signature of an optimized orthosteric pocket. In the Dasatinib complex, the ATP-binding cleft (residues 315–321) is dominated by minimally frustrated residues, forming a stable, low-frustration interaction core. Concurrently, discrete clusters of high frustration are localized in regulatory regions, including the C-lobe (residues 320–340), the A-loop and P+1 loop (residues 395–410), and the αG-helix (residues 440–455). These high-frustration zones correspond to conformationally adaptable motifs that are implicated in functional switching (Figure 5-figure supplement 1). A nearly identical frustration pattern is observed in the Axitinib complex, where the orthosteric site is similarly enriched in minimal frustration, consistent with its conserved hydrogen-bonding network (e.g., with T315, K271, Y253, and F382) (Figure 5-figure supplement 2). Notably, the mutational frustration profiles for both inhibitors show an even stronger concentration of minimal frustration in the binding site than the conformational profiles, highlighting the evolutionary conservation and sequence constraint characteristic of orthosteric pockets. These results directly exemplify the “energetically locked” architecture described for type I inhibitors in the main text: the orthosteric site is a hotspot of minimal frustration reflecting evolutionary optimization and structural invariance, whereas surrounding regulatory elements exhibit elevated frustration, enabling the conformational plasticity required for allosteric regulation and state transitions.

### Dynamic Frustration Signatures of the ABL Inactive States : Persistent Neutrality of Myristoyl Pocket and Shift to High Frustration in Regulatory Sites Mediating Transitions

In contrast, allosteric regulation introduces a distinct energetic signature characterized by persistent neutral frustration (NF). Even in the maximally stabilized, autoinhibited assembly bound to both Nilotinib and Asciminib (Figure 5-figure supplement 3), the allosteric myristoyl pocket remains heavily neutrally frustrated, suggesting this region is evolutionarily ‘concealed’ to maintain functional plasticity. This persistent NF signature is similarly observed in the kinase domain when bound to the foundational allosteric inhibitor GNF-2 (Figure 5-figure supplement 4). Notably, this neutral landscape is not restricted to inhibited states; the allosteric activator DPH (Figure 5-figure supplement 5) also targets a region of high NF density, further confirming that these regulatory pockets lack the strong coevolutionary optimization seen at orthosteric sites. Importantly, these complexes consistently display a predominance of neutrally frustrated residues enriched in and around the allosteric myristoyl pocket. This pattern contrasts sharply with the minimal-frustration signature of orthosteric sites and highlights the evolutionary plasticity and structural adaptability of allosteric regions. Despite the overall prevalence of neutral frustration, we observed appreciable redistribution from neutrally frustrated to highly frustrated clusters in in key regulatory junctions implicated in functional kinase transitions (Figure 5-figure supplement 3-5). Overall, the frustration signatures of allosteric ABL complexes illustrate an adaptive energetic regime: a backbone of neutral frustration that permits conformational flexibility, interspersed with accumulation of high-frustration clusters at regulatory interfaces in inactive states, creating a driving force for conformational switching.

The important finding of this analysis is seemingly persistent neutrality of the allosteric myristoyl pocket across different ABL states and complexes. Although Asciminib, GNF-2, and an allosteric activator DPH occupy the same myristoyl pocket, each induces a distinct local conformational response, demonstrating that the pocket is not a fixed cavity but a deformable regulatory interface whose geometry is reshaped by ligand-dependent redistribution of frustration. Upon Asciminib binding, the αI-helix undergoes a pronounced bending motion that stabilizes the autoinhibited state (Figure 5E). The residues accommodating this deformation are largely neutrally frustrated, with embedded highly frustrated positions acting as mechanical pivots that enable strain redistribution without enforcing strict sequence conservation. Binding of GNF-2 produces a different deformation of the same helix and surrounding regulatory elements, yielding an alternative inactive configuration (Figure 5F). Compared with Asciminib, the magnitude and direction of helix bending differ, and adjacent loops and spine elements reorganize in a ligand-specific manner. In contrast, binding of the allosteric activator DPH partially reverses this deformation, unbending the αI-helix and promoting a more active-like configuration (Figure 5G). This inversion of response-activation rather than inhibition-demonstrates that the same pocket can support opposing functional outcomes. These differences show that the pocket does not encode a single canonical inhibitory geometry but supports multiple low-energy conformations depending on how local frustration is relieved.

Combined, these structural responses of the ABL allosteric sites to binding of different modulators exemplifies the defining feature of the allosteric sites across human kinome characterized by context-dependent remodeling. Neutral frustration of the energy landscapes for the allosteric binding regions ensures mutational tolerance and conformational diversity, while localized high frustration enables sensitivity to perturbation and signal propagation. The ABL analysis provides atomic-scale validation of the global frustration–predictability relationship. Orthosteric sites are encoded as energetically optimized basins, producing conserved sequence and geometric patterns that PLMs and structure-based methods detect with high confidence. By contrast, allosteric sites are encoded as energetically permissive and heterogeneous regions whose defining features are conformational strain and context-dependent remodeling rather than conserved motifs. The invisibility of allosteric pockets to AI predictors is not accidental but emerges from their energetic architecture: neutral frustration masks evolutionary regularities while localized high frustration encodes regulatory sensitivity.

### Conserved Island of Neutrality: Neutral Frustration of ABL Myristoyl Allosteric Pocket Persists Across Distinct Allosteric Modulators and Regulatory Domains

To investigate how neutral frustration behaves under complex regulatory conditions that more closely mirror the cellular context, we performed comparative frustration mapping on three distinct structural states of the ABL kinase that include the SH2-SH3 regulatory domains (Figure 6). This series captures the ABL kinase across key regulatory states-from the unliganded autoinhibited assembly through the myristoyl-stabilized inactive conformation to inhibitor-bound forms-providing a window into how frustration signatures respond to both protein-protein interactions and small-molecule binding. We first examined isolated ABL kinase domain complexes bound to chemically and functionally diverse allosteric modulators (Figure 6A–D). Despite substantial differences in ligand chemistry and functional outcomes, the myristoyl pocket consistently displays high-density neutral frustration (blue patches) across all complexes. In the ABL-GNF-2 complex (PDB 3K5V, Figure 6A), the allosteric inhibitor GNF-2 binds the myristoyl pocket while Imatinib occupies the orthosteric ATP cleft. The pocket-lining residues exhibit prominent neutral frustration density, reflecting the conformational plasticity that enables GNF-2 to stabilize the autoinhibited state through αI-helix bending. Similarly, in the ABL-DPH complex (PDB 3PYY, Figure 6B), the allosteric activator DPH binds the same myristoyl pocket yet produces the opposite functional outcome-promoting an active-like conformation through αI-helix unbending. Despite this functional inversion, the pocket retains its neutral frustration signature, indicating that the underlying energetic architecture permits both inhibitory and activating conformational responses.

**Figure 6.**
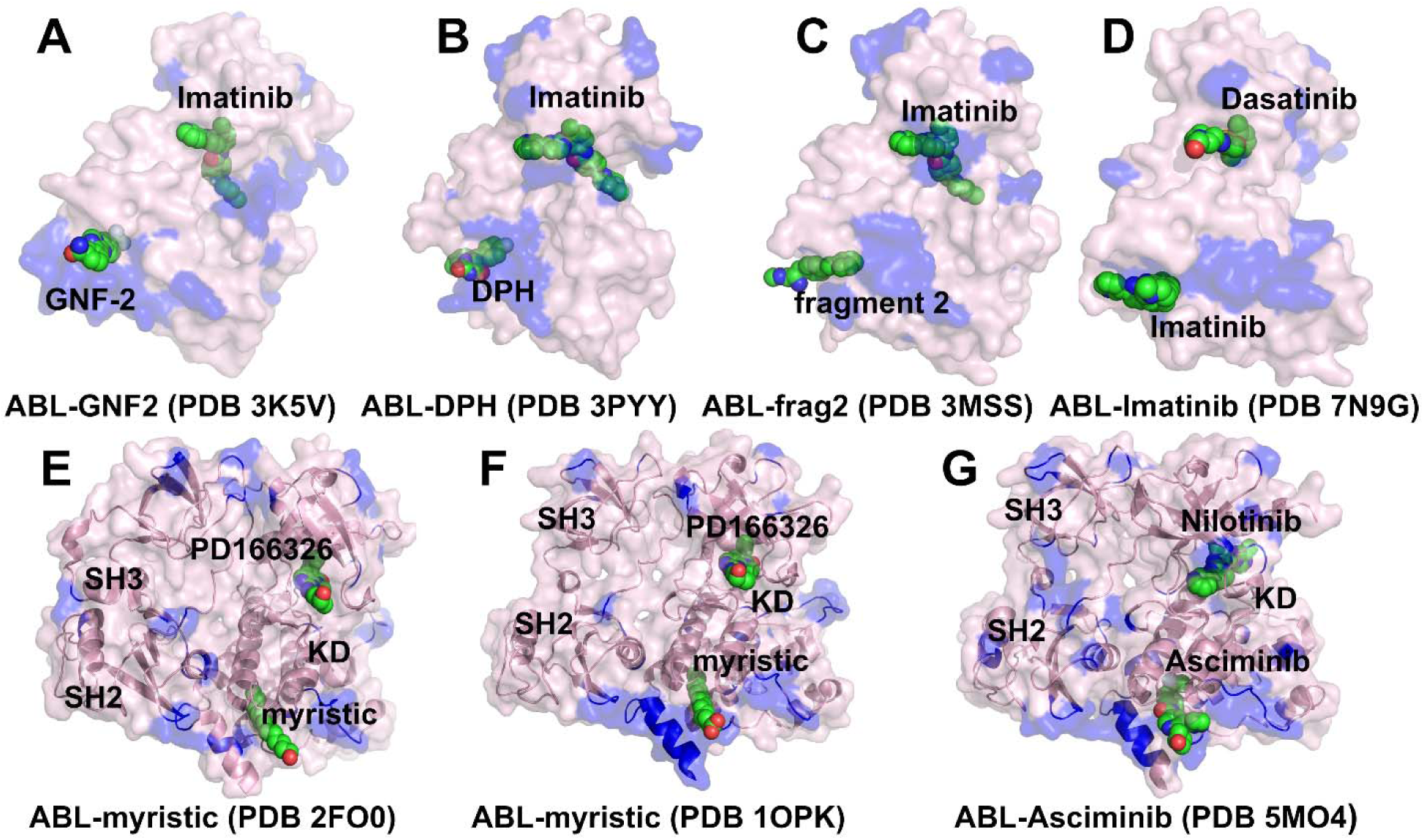
Structural diversity and persistent neutral frustration signatures at the ABL myristoyl allosteric pocket across regulatory states. (A–D) Isolated kinase domain complexes with diverse allosteric modulators. Surface representations of the ABL kinase domain bound to distinct ligands in the orthosteric and allosteric sites. Blue patches indicate regions of high-density neutral frustration. (A) ABL bound to the allosteric inhibitor GNF-2 (green spheres) in the myristoyl pocket and Imatinib (orthosteric) in the ATP cleft (PDB 3K5V). (B) ABL bound to the allosteric activator DPH (green/red spheres) in the myristoyl pocket and Imatinib (orthosteric) (PDB 3PYY). (C) ABL bound to allosteric fragment 2 (green spheres) and Imatinib (orthosteric) (PDB 3MSS). (D) ABL bound to Imatinib (green spheres) in the allosteric pocket and Dasatinib (orthosteric) in the ATP cleft, illustrating allosteric binding of an ATP-competitive inhibitor in resistant variants (PDB 7N9G). In all cases, the allosteric pocket is lined by neutral frustration hotspots (blue), reflecting conformational plasticity. (E–G) Full SH3–SH2–Kinase Domain (KD) regulatory assemblies. Ribbon and surface representations of the autoinhibited ABL assembly, showing the SH3 (dark blue), SH2 (light blue), and Kinase Domain (pink/white) domains. Blue patches on the kinase domain surface indicate neutral frustration density. (E) Autoinhibited assembly with myristic acid (green/red spheres) bound in the allosteric pocket and PD166326 (orthosteric) (PDB 2FO0). (F) Autoinhibited assembly with myristic acid bound in the allosteric pocket and PD166326 (orthosteric) (PDB 1OPK). (G) Autoinhibited assembly with the allosteric inhibitor Asciminib (green/red spheres) bound in the myristoyl pocket and Nilotinib (orthosteric) (PDB 5MO4).

The ABL-fragment 2 complex^85^ (PDB 3MSS, Figure 6C) demonstrates that even small molecular promotes an extended active-like conformation and exerts dual negative effects on inhibition, fragments binding the myristoyl pocket encounter the same neutrally frustrated environment. Most strikingly, in the ABL-Imatinib complex (PDB 7N9G, Figure 6D), Imatinib-originally designed as an ATP-competitive inhibitor-binds the allosteric myristoyl pocket in drug-resistant variants, while Dasatinib occupies the orthosteric site. This unexpected allosteric binding mode, which occurs within the same neutrally frustrated pocket architecture observed with dedicated allosteric modulators. The critical observation across all four complexes is that the allosteric pocket is consistently lined by neutral frustration hotspots (blue patches), regardless of ligand identity or functional outcome. This persistence indicates that neutral frustration is not a consequence of specific ligand chemistry but an intrinsic property of the pocket that enables its remarkable regulatory versatility. The neutrally frustrated environment permits the αI-helix to sample multiple conformations-bent (GNF-2, fragment 2), partially unbent (DPH), or extended (Imatinib in resistant variants) without requiring remodeling of the underlying energy landscape. We argue that allosteric modulators merely select among pre-existing conformational states encoded by neutral frustration rather than inducing new energetic minima.

To determine whether neutral frustration persists in more physiologically relevant regulatory contexts, we examined full SH3–SH2–Kinase Domain (KD) autoinhibited assemblies (Figure 6E–G). These complexes include the upstream regulatory domains that stabilize the autoinhibited state through interdomain contacts, providing a more complete picture of how the myristoyl pocket functions within the native regulatory architecture. In the apo SH3-SH2-KD assembly (PDB 2FO0, Figure 6E), the myristoyl pocket is occupied by endogenous myristic acid while PD166326 binds the orthosteric site. The SH3 (dark blue) and SH2 (light blue) domains clamp the kinase domain (pink/white) in the autoinhibited conformation. Despite this extensive interdomain stabilization, the myristoyl pocket retains its characteristic neutral frustration density (blue patches), extending into the adjacent αI-helix that mechanically couples the pocket to the SH2-kinase interface. The same pattern holds in the SH3-SH2-KD assembly with myristic acid and PD166326 (PDB 1OPK, Figure 6F), where the regulatory domains and orthosteric inhibitor jointly stabilize the inactive state. The myristoyl pocket remains neutrally frustrated, indicating that even when the pocket is occupied by its physiological ligand and the kinase is fully autoinhibited, the pocket retains conformational plasticity rather than settling into a single rigid minimum. In the SH3-SH2-KD-Asciminib-Nilotinib complex (PDB 5MO4, Figure 6G), the clinically approved allosteric inhibitor Asciminib binds the myristoyl pocket while Nilotinib occupies the orthosteric site, with the SH3-SH2 clamp maintaining autoinhibition. Despite simultaneous engagement of both orthosteric and allosteric sites by clinical inhibitors, and despite the stabilizing influence of the SH3-SH2 regulatory domains, the myristoyl pocket and αI-helix retain their neutral frustration signature.

The persistence of neutral frustration across all three regulatory assemblies carries profound biological implications. The SH3-SH2 clamp provides a stable “down-regulatory” anchor that rigidifies the core and suppresses catalytic activity, yet it leaves the allosteric pocket poised for modulation. The regulatory domains and the allosteric pocket thus operate on different levels: the domains set the overall regulatory context through interdomain contacts, while the pocket’s neutral frustration ensures continued responsiveness to diverse inputs-whether endogenous myristoyl insertion, allosteric drug binding, or mutations that alter pocket affinity.

Taken together, the examined seven complexes establish the ABL myristoyl pocket as a paradigmatic example of intrinsically encoded and persistent neutral frustration. Across isolated kinase domains and full regulatory assemblies, across diverse ligands (GNF-2, DPH, fragment 2, Imatinib, myristic acid, Asciminib), and across functional outcomes (inhibition, activation, resistance), the pocket maintains its neutral frustration signature. The allosteric pocket is integrated into broader allosteric networks that connect this distal site to the ATP-binding cleft, ensuring that local binding events propagate across the entire structure. This architectural design enables the pocket to serve its biological function: remaining continuously available for context-dependent modulation while providing a druggable target for allosteric inhibitors. The ABL analysis provides atomic-resolution validation of the principles inferred from kinome-wide data. The enrichment of minimal frustration at orthosteric sites, the dominance of neutral frustration at allosteric sites, the spatial segregation of frustrated regions from the catalytic core, and the persistence of neutral frustration across diverse regulatory states-all of these findings mirror and reinforce the patterns observed across the broader kinase family.

## Discussion

The systematic use of AI models as diagnostic probes across the human kinome reveals a central principle: binding-site detectability reflects the underlying biophysical organization of protein energy landscapes rather than algorithmic limitations. By repurposing PLMs from predictive tools into instruments for interrogating evolutionary design, we show that these models are intrinsically sensitive to minimally frustrated regions-structurally constrained, catalytically essential cores shaped by strong purifying selection. In contrast, allosteric sites consistently localize within neutrally frustrated environments that enable conformational plasticity and mutational tolerance. This evolutionary permissiveness weakens sequence-level constraints and erodes the statistical signals required for inference, giving rise to an intrinsic “allosteric blind spot.”

A key advance of this work is the introduction of local frustration analysis as an interpretability layer that links machine learning outputs to physical principles. The strong correspondence between PLM prediction patterns and frustration classes provides a mechanistic explanation for why current AI models succeed in identifying orthosteric sites yet struggle with allosteric pockets. Orthosteric sites reside in deep, funneled energetic basins that generate robust coevolutionary signals, whereas allosteric sites occupy shallow, rugged landscapes where functional adaptability is achieved at the expense of sequence-based detectability. In this context, model uncertainty is not noise but a meaningful signature of regulatory potential.

Protein kinases provide a uniquely well-controlled system for uncovering this relationship. The KinCoRe framework offers high-resolution structural annotation across 453 human kinases, spanning multiple conformational states and five mechanistically distinct inhibitor classes. This extensive experimental foundation ensures that the observed dichotomy between orthosteric and allosteric detection arises from genuine energetic and evolutionary principles rather than dataset bias. The availability of diverse structural ensembles further enables frustration analysis to capture the energetic determinants of conformational transitions central to kinase regulation.

Importantly, our findings align with independent experimental evidence from deep mutational scanning studies, which show that allosteric regulation is widespread, distributed, and frequently mediated by distal, surface-exposed regions. These observations can now be interpreted through the lens of neutral frustration: sites that tolerate mutation while modulating function correspond to energetically permissive regions that support evolutionary exploration without destabilizing the protein core. This convergence between experiment and computation elevates neutral frustration from a theoretical construct to a general organizing principle of allosteric regulation.

The landscape-based frustration analysis provides a principled basis for prioritizing experimental validation, bridging the gap between computational prediction and mechanistic discovery. Current models implicitly assume that functional relevance is encoded in conserved patterns-an assumption that holds for orthosteric sites but fails for allosteric regions where regulatory function depends on conformational plasticity. Overcoming this intrinsic blind spot requires incorporating energy landscape signatures directly into learning objectives. Frustration indices may serve as natural priors: as regularization terms penalizing overconfidence in neutrally frustrated zones, as edge weights in graph neural networks to capture allosteric couplings, or as targets in multi-task learning frameworks that force models to internalize the relationship between evolutionary signal and energetic organization. This study highlights the importance of aligning AI approaches with the physical principles governing biomolecular systems, providing a foundation for the development of interpretable and mechanistically grounded AI frameworks.

## Conclusions

The central finding of this work is that the detectability of protein binding sites by machine learning models is fundamentally governed by the biophysical character of the target site-a principle that emerges from integrating PLMs as diagnostic probes with energy landscape theory. Across a rigorously curated dataset of human kinase–ligand complexes, we demonstrate a sharp and reproducible dichotomy: orthosteric ATP-binding sites are detected with high precision by sequence-based models, while allosteric sites evade detection despite preserved ranking ability. Critically, this performance gap is not algorithmically driven but represents a faithful readout of the underlying energy landscape organization.

Local frustration analysis reveals the physical basis for this dichotomy. Interpreting these patterns through local frustration analysis reveals that model “visibility” is closely linked to energetic structure. Minimally frustrated regions encode stable, evolutionarily conserved interactions that give rise to strong sequence signals, while neutrally frustrated regions support conformational flexibility and regulatory diversity but lack consistent evolutionary signatures. In this context, prediction confidence can be understood as a readout of how strongly functional constraints are embedded in sequence space. Rather than viewing these differences as model deficiencies, our results position them as diagnostic signals that expose the biophysical logic of protein regulation.

This perspective reframes explainable AI as a tool for probing the relationship between sequence, structure, and energetics, enabling systematic identification of regions where functional roles are not directly encoded in evolutionary conservation. By demonstrating that algorithmic visibility is governed by energetic frustration, we establish a quantitative baseline against which future physics-informed models can be calibrated. As high-quality structural and mutational data continue to accumulate across protein families, the principles elucidated here may inform the development of interpretable, mechanistically grounded AI tools. More broadly, this work outlines an explainable AI strategy for bridging data-driven learning with energy landscape theory, transforming predictive models into tools for probing biological phenomena and mechanistic inference enabling a deeper understanding of how evolution encodes regulatory complexity and allostery in proteins.

## Materials and Methods

### Dataset Curation and Evaluation Framework

To ensure rigorous evaluation of model generalization and avoid potential train–test leakage, we implemented a sequence similarity filtering strategy between training and evaluation datasets. The protein language model was trained on a subset of the LIGYSIS dataset (3,006 sequences).

To eliminate overlap with the kinase dataset used for evaluation, all training sequences sharing greater than 30% sequence identity with any protein in the KinCoRe dataset were removed using MMseqs2 clustering.^86–88^ No representative selection was performed; any sequence exceeding the similarity threshold was excluded. This filtering resulted in a reduced training set of 2,949 sequences (denoted LIGYSIS_SI30), ensuring that all reported results reflect genuine generalization rather than memorization. The evaluation dataset was derived from the KinCoRe database^20–22^, yielding a total of 9,901 kinase–ligand complexes (structure–chain pairs) spanning 453 human kinases (Table 1-Source Data 1). The dataset includes five mechanistic classes of inhibitor binding defined by DFG motif conformation and αC-helix position: Type I, Type I.5, Type II (orthosteric), and Type III and Type IV (allosteric). Because structural datasets are inherently imbalanced (Table 1-Source Data 2), with certain kinases heavily overrepresented we constructed a non-redundant subset (KinCoRe-UNQ)^68^ by selecting one structure per UniProt accession (708 complexes). This controls for biases arising from repeated structures of identical or highly similar sequences. Comparative analysis between the full dataset and KinCoRe-UNQ demonstrated minimal differences in model performance, indicating that results are not driven by redundancy. Importantly, the dataset exhibits a strong class imbalance, with orthosteric sites (Types I, I.5, II) comprising ∼90% of all complexes, while allosteric sites (Types III and IV) account for only ∼7%.

**Table 1.** Composition of the kinase dataset derived from KinCoRe.

| Inhibitor class | Binding mode | KinCoRe (full) | KinCoRe-UNQ |
| --- | --- | --- | --- |
| **Type I** | ATP-competitive (DFG-in, C-in) | 7,604 | 407 |
| **Type I.5** | ATP-competitive (C-out extension) | 775 | 117 |
| **Type II** | ATP-competitive (DFG-out) | 553 | 65 |
| **Type III** | Proximal allosteric | 294 | 30 |
| **Type IV (ALLO)** | Distal allosteric | 675 | 89 |
| **Total** | - | 9,901 | 708 |

**Table 1 Source Data 1.** Grouped categories used for PLM performance benchmarking.

| Site category | Classes included | Count | Fraction (%) |
| --- | --- | --- | --- |
| **Orthosteric** | Type I, I.5, II | 8,932 | 90.2% |
| **Allosteric (proximal)** | Type III | 294 | 3.0% |
| **Allosteric (distal / ALLO)** | Type IV | 675 | 6.8% |

**Table 1 Source Data 2.**
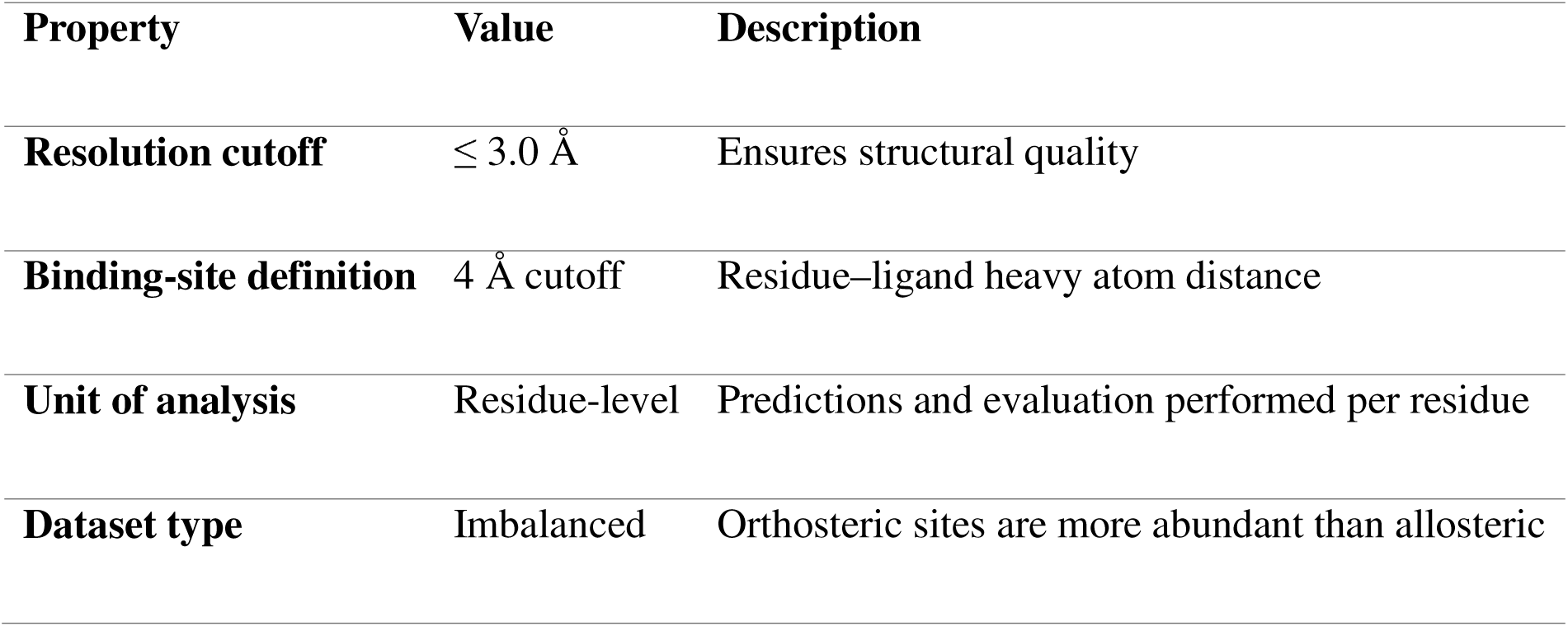
Structural and data characteristics.

### Protein Representation and Model Architecture

Protein sequences were encoded using the ESM-2 (650M) protein language model^27^, a transformer-based architecture trained on large-scale sequence data to capture evolutionary and contextual relationships. In this work, the model is not treated solely as a predictive tool, but as a diagnostic probe of sequence-encoded biophysical information, where prediction performance reflects the degree to which functional sites are evolutionarily constrained and therefore “visible” in sequence space. For each sequence, residue-level embeddings (1,280 dimensions) were extracted and used as input features. These embeddings encode both local physicochemical context and long-range sequence dependencies, enabling structure-aware inference without explicit structural input. In this framework, PLMs are not treated as end-point predictors but as diagnostic probes of sequence-encoded biophysical information. A lightweight classification head was applied to map residue embeddings to binding-site probabilities.^68^ This design preserves the underlying evolutionary representation while enabling task-specific interpretation. We implemented a two-stage hierarchical fine-tuning protocol: In Phase I (head-only training), the transformer encoder was frozen, and only the classification head parameters were optimized for two epochs. This establishes a stable mapping between embeddings and binding-site annotations without perturbing pre-trained representations. In Phase II (end-to-end fine-tuning), all model parameters were unfrozen and jointly optimized for one additional epoch, allowing adaptation to kinase-specific patterns while maintaining global sequence features. This two-stage approach proved essential for achieving robust performance across diverse kinase families. Preliminary experiments with immediate end-to-end fine-tuning resulted in unstable convergence and reduced generalization, particularly for allosteric sites with weak evolutionary signals.^68^

### Prediction and Evaluation Framework

The model outputs residue-level probabilities *p*_i_ E [0,1], which were evaluated using both threshold-free and threshold-dependent metrics. Threshold-free performance was assessed using Area under the receiver operating characteristic curve (AUROC) and Area under the precision–recall curve (AUPR). For binary classification, two thresholding strategies were used: Fixed threshold (PLM-fixed) with a global threshold (t = 0.75) selected by maximizing Matthews correlation coefficient (MCC) on validation data and class-optimized threshold (PLM-bestMCC) where thresholds were optimized separately for each inhibitor class via grid search (t ∈ [0,1], step 0.05), accounting for differences in evolutionary constraint between orthosteric and allosteric sites. Binary predictions were evaluated using MCC, F1 score, and accuracy. Analyses were performed at both micro-averaged (global) and macro-averaged (per-structure) levels to account for class imbalance and structural heterogeneity.

### Structural Analysis and Preparation of Kinase Structures

To establish a direct correspondence between sequence-based predictions and the physical organization of kinase structures, we constructed a structurally standardized ensemble of kinase domains spanning major families, conformational states, and inhibitor classes represented in the KinCoRe dataset. This step was essential to ensure that both prediction outputs and energetic analyses are interpreted within a consistent structural and biophysical framework. Representative structures were selected from the Protein Data Bank based on stringent criteria, including high crystallographic resolution (≤3.0 Å), coverage of major kinase groups (AGC, CAMK, CMGC, STE, TK, and TKL), and representation of diverse inhibitor binding modes (Types I–IV). Selection was guided to preserve the conformational diversity captured by KinCoRe, including active (DFG-in/αC-in) and inactive (DFG-out or αC-out) states, thereby enabling systematic comparison across orthosteric and allosteric regulatory regimes.

All structures underwent a uniform and rigorous preprocessing pipeline to ensure compatibility with downstream energetic analysis. This included removal of crystallographic artifacts (water molecules, buffer components, and non-cognate ligands), retention of the biologically relevant inhibitor where appropriate, and reconstruction of missing side chains and unresolved local regions when feasible. All structures were obtained from the Protein Data Bank.^89^ Hydrogen atoms and missing residues were initially added and assigned according to the WHATIF program web interface.^90,91^ The structures were further pre-processed through the Protein Preparation Wizard (Schrödinger, LLC, New York, NY) for assignment and adjustment of ionization states, formation of assignment of partial charges as well as additional check for possible missing atoms and side chains that were not assigned by the WHATIF program. The missing loops in the cryo-EM structures were reconstructed using template-based loop prediction approaches ModLoop^92^ and ArchPRED^93^ and further confirmed by FALC (Fragment Assembly and Loop Closure) program.^94^ The side chain rotamers were refined and optimized by SCWRL4 tool.^95^

Prior to frustration analysis, all kinase-ligand complexes were subjected to a multi-stage structural refinement and energy minimization protocol to resolve steric clashes and optimize hydrogen-bonding networks. The AMBER22 suite was used for all refinements.^96^ Protein atoms were modeled using the ff19SB force field, while ligand parameters were generated via the General Amber Force Field (GAFF2)^97^ with AMBER antechamber module.^97,98^ To prevent structural distortion, a layered minimization protocol was executed: (a) Hydrogen-Only Relaxation: 1,000 steps of steepest descent were performed with all heavy atoms (protein and ligand) restrained, allowing for the optimization of hydrogen atom positions and protonation states; (b) Solvent and Ligand Minimization: 2,000 steps focusing on the ligand and surrounding water molecules, with the protein backbone and side chains held under harmonic restraints. (c) Side Chain Relaxation: 2,000 steps targeting protein side chains to resolve localized atomic overlaps within the protein interior; (d) Backbone Minimization: 2,000 steps of backbone-only refinement to ensure secondary structure stability. (e) Unrestrained All-Atom Refinement: A final 5,000-step gradient minimization was performed with all restraints removed. This ensured the complex reached a local potential energy minimum, providing a consistent energetic baseline for the subsequent calculation of configurational and mutational frustration indices. To enable cross-structure comparison, kinase domains were structurally aligned using conserved elements of the catalytic core. Specifically, alignment was performed based on Cα atoms of the central β-sheet and key conserved motifs, including the HRD catalytic loop and the DFG motif. This procedure establishes a common structural reference frame that preserves functional architecture while minimizing variability arising from peripheral regions.

This structurally harmonized ensemble provides the foundation for two key analyses. First, it enables consistent mapping of residue-level prediction probabilities onto three-dimensional structures, allowing direct visualization of model-detected binding regions across kinase families. Second, it ensures that local frustration calculations are performed on energetically comparable and physically consistent conformations, which is critical for interpreting frustration signatures in terms of intrinsic energy landscape features rather than structural artifacts.

By integrating KinCoRe classification with rigorous structural normalization, this framework establishes a unified basis for linking sequence-derived predictions, structural organization, and energetic signatures, enabling a physically grounded interpretation of orthosteric and allosteric site architectures.

### Local Frustration Analysis

To characterize the energetically encoded signatures of functional sites, we employed a local frustration analysis framework rooted in energy landscape theory.^4–42^ This approach quantifies the extent to which local interactions within the native protein structure are energetically optimized or strained, distinguishing between residues that are evolutionarily and structurally stabilized from those involved in conformational adaptability or functional stress. Two complementary frustration indices were computed. Configurational (or conformational) frustration assesses the sensitivity of a native contact to local structural perturbations. It is calculated by randomizing both residue identities and interatomic distances within the native contact geometry, thereby measuring whether the observed interaction geometry is more favorable than structurally plausible alternatives. Mutational frustration evaluates the energetic optimality of a given residue–residue pair by comparing its interaction energy to that of all possible amino acid substitutions at the same positions, while the backbone conformation remains fixed. This metric captures evolutionary constraints by reflecting whether the native amino acid pairing is more favorable than evolutionary accessible alternatives.^42,43^ Both frustration indices were expressed as Z-scores, defined as:

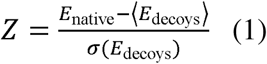

where *E*_native_is the energy of the native contact, and 〈*E*_decoys_〉 and *σ* (*E*_decoys_)represent the mean and standard deviation, respectively, of energies computed from an ensemble of 1,000 structural decoys generated by perturbing local conformations or residue identities.

Following well-established thresholds validated across diverse protein systems^42,43^, contacts were classified as minimally frustrated (Z > 0.78). In this regime, the native interaction is significantly more favorable than decoy alternatives, indicating strong energetic optimization. Highly frustrated (Z < −1.0). In the high frustration regime, the native interaction is significantly less favorable, reflecting local strain or conflict. Neutrally frustrated (−1.0 ≤ Z ≤ 0.78) which is a regime when the native interaction is neither strongly favored nor disfavored, consistent with conformational plasticity or mutational tolerance. To map these interactions onto a residue-level profile, the local density of frustrated contacts within a 5 Å radius of each residue was computed. A residue was assigned a dominant frustration category if more than 50% of its interacting contacts within the most populated conformational ensemble belonged to the same frustration class.

### Statistical Analysis

Statistical significance of performance differences between the PLM and baseline methods was determined using paired two-tailed t-tests, with p < 0.05 considered significant. For multi-family comparisons (e.g., across five kinase types), p-values were adjusted via the Benjamini–Hochberg false discovery rate (FDR) correction.^99^ To ensure a balanced interpretation under class-imbalanced conditions, both micro- and macro-averaging schemes were employed. Micro-averaged metrics were computed globally across all residues, while macro-averaged metrics were calculated per structure and then averaged across the dataset, giving equal weight to each kinase irrespective of chain length or residue count. For the frustration analysis, residue-level frustration densities were compared across orthosteric and allosteric sites using the Mann–Whitney U test, a non-parametric method appropriate for non-normal distributions and unequal variances.^100^ All statistical computations were performed in Python 3.10 using SciPy (v1.10) for hypothesis testing, statsmodels (v0.14) for multiple comparison correction, and custom scripts for effect size and correlation analysis. Visualization of distributions and significance annotations were generated with Matplotlib (v3.7) and Seaborn (v0.12).

## Author Contributions

Conceptualization, G.V.; Methodology, W.G., M.L., L.T., B.F., K.R., V.S., M.N., D.H., G.V.; Software, W.G., M.L., L.T., B.F., K.R., V.S. M.N., D.H., G.V.; Validation, W.G., M.L., L.T., B.F., K.R., V.S., D.H., G.V.; Formal analysis, W.G., M.L., L.T., B.F., K.R., V.S., D.H., G.V.; Resources, W.G., M.L., L.T., B.F., K.R., V.S., D.H., G.V.; Data curation, W.G., M.L., L.T., B.F., K.R., V.S., D.H., G.V.; Writing-original draft preparation, K.R., V.S., D.H., G.V. ; Writing-review and editing, K.R., V.S., D.H., G.V. ; Visualization. W.G., M.L., L.T., B.F., K.R., V.S., M.N., D.H., G.V.; Supervision D.H., G.V.; Project administration, D.H., G.V. ;. Funding acquisition, D.H., G.V.

All authors have read and agreed to their individual contributions. All authors have approved the submission of this manuscript.

## Data Availability Statement

Data for this article, including description of data types, all data, software and scripts are freely available at the Github repository https://github.com/kiarka7/plm-vs-p2rank-kinases and the Zenodo archive (https://doi.org/10.5281/zenodo.18226590).

The fine-tuned PLM model can be downloaded from this data storage: https://owncloud.cesnet.cz/index.php/s/8RSJqt60D2uWJNa

The code and datasets related to this PLM are available at: https://github.com/skrhakv/LBS-pLM

The Github repository https://github.com/kiarka7/plm-vs-p2rank-kinases contains publicly available scripts required to reproduce both PLM and P2Rank predictions reported in the study.

Inference was performed using custom Python scripts that wrap the fine-tuned ESM2 checkpoint and implement residue extraction, tokenization, batching, and model evaluation. All scripts required to reproduce the PLM predictions reported here are publicly available in GitHub repository (https://github.com/kiarka7/plm-vs-p2rank-kinases). The fine-tuned model checkpoint can be provided upon request for reproducibility.

Zenodo archive (https://doi.org/10.5281/zenodo.18226590) provides the datasets, processed prediction outputs, evaluation results, figures, and scripts used in the study. The archive mirrors the directory structure expected by the analysis pipeline and contains two dataset variants: (a) KinCoRe, which includes all protein–ligand complexes used for the main analyses reported in the manuscript and (b) KinCoRe-UNQ, a reduced variant retaining a single representative structure per protein (UniProt ID), which was used for Supporting Information analyses to control for over-representation of proteins with multiple solved structures. All results presented in the manuscript are derived from the KinCoRe dataset, while KinCoRe-UNQ is used for analyses reported in the Supporting Information. Protein structures are not included in this archive. Crystal structures were obtained and downloaded from the Protein Data Bank (http://www.rcsb.org). The rendering of protein structures was done with UCSF ChimeraX package (https://www.rbvi.ucsf.edu/chimerax/) and Pymol (https://pymol.org/2/) . All structures are publicly available from the RCSB Protein Data Bank and can be automatically downloaded using the provided scripts.

## Acknowledgments

This research was funded by the National Institutes of Health under Award 1R01AI181600-01, 5R01AI181600-02 and Subaward 6069-SC24-11 to G.V. The authors acknowledge the use of high-performance computing (HPC) resources and computational infrastructure provided by Schmid College of Science and Technology and the Keck Center for Science and Engineering at Chapman University. G.V. acknowledges support from Schmid College of Science and Technology for providing the computing resources necessary for this study.

