## Supplemental Materials: Figure Supplements 1-1, 2-1, 2-2, 4-1, 5-1, 5-2, 5-3, 5-4, 5-5 and Supplemental Tables S1-S5 (Figure 3-source data 1). for "Decoding Allosteric Grammar with Explainable AI Integrating Protein Language Models and Energy Landscape Analysis: Neutral Frustration at Allosteric Binding Sites Encodes Regulatory Versatility in Protein Kinases": SUPPLEMENTAL_MATERIALS_BIORXIV.pdf

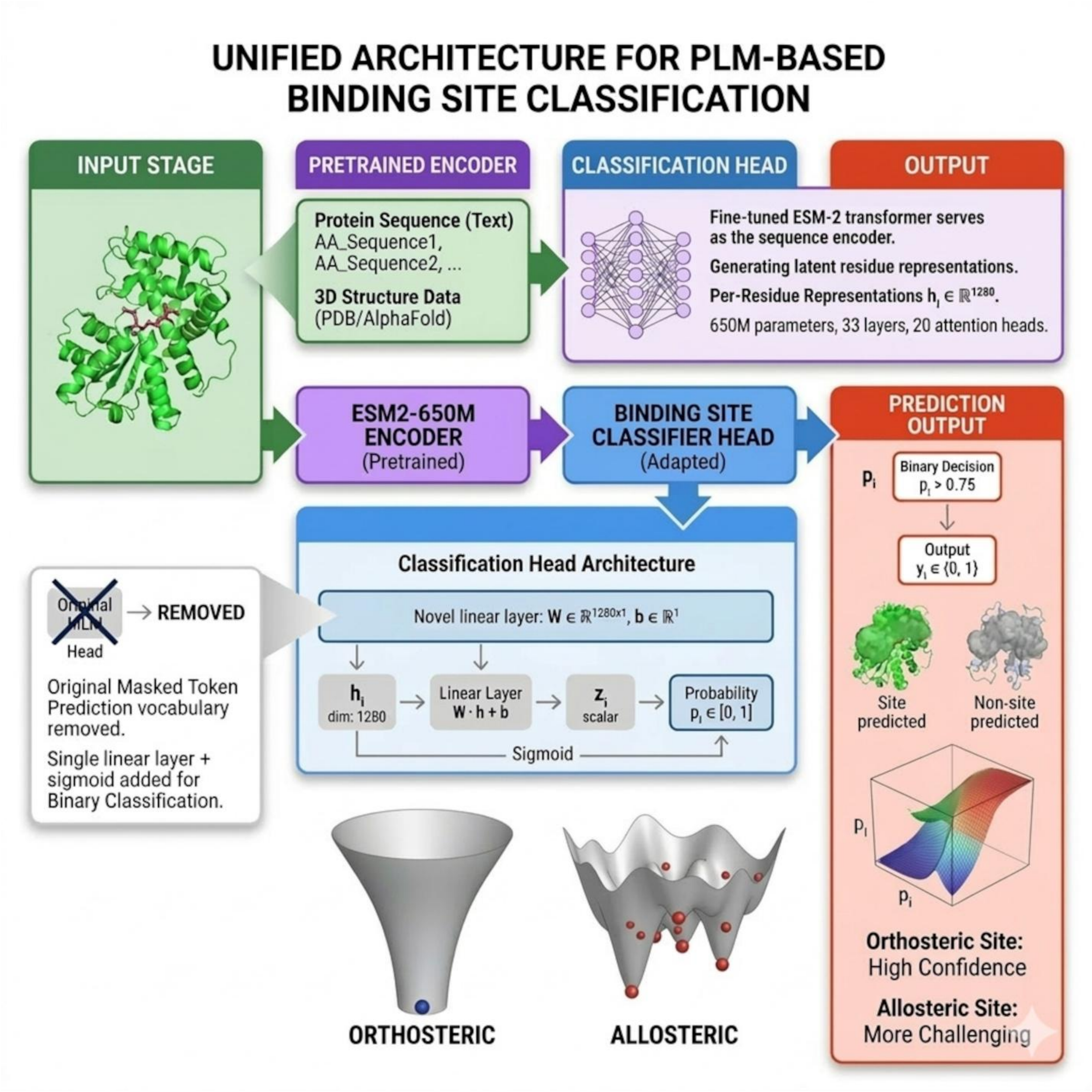


**Figure 1-figure supplement 1. Schematic of the unified PLM-based architecture and residue-level classification pipeline.** Detailed representation of the computational stages used to map sequence and structural data to binding site predictions. Input and Encoding Stage (Top Panel): Protein primary sequences and 3D coordinates (from PDB or AlphaFold) are processed by a 33-layer ESM-2 transformer encoder[27] (650M parameters, 20 attention heads). The encoder generates 1280-dimensional latent representations (hi) for every residue, capturing both local and distal evolutionary context. Classification Head Adaptation (Center Panel): The original Masked Language Model (MLM) head is replaced with a specialized linear layer (W∈R1280×1) and a sigmoid function. This architecture converts the latent residue vectors into a scalar probability (pi ∈[0,1]) for binary site classification. Prediction Output and Conceptual Landscapes (Bottom Panel): Residues meeting the classification threshold (pi>0.75) are mapped to the 3D structure. The schematic energy landscapes illustrate the underlying difficulty of the task: orthosteric sites occupy deep, stable basins that yield high-confidence predictions, whereas allosteric sites-defined by rugged, multi-minima landscapes-present more challenging targets for the sequence-based model.

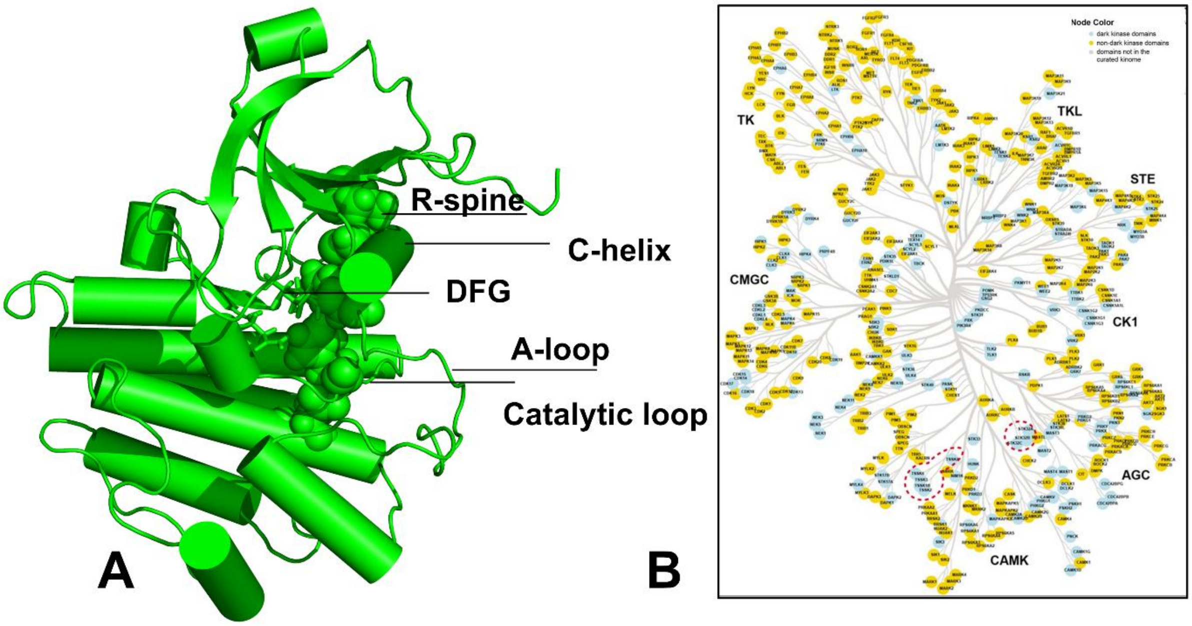

**Figure 2-figure supplement 1. Structural motifs of the kinase catalytic domain and kinome-wide distribution of study targets.** (A) Representative 3D structural model of the conserved kinase fold highlighting the regulatory architecture. The Regulatory spine (R-spine) is shown in a vertically aligned, "assembled" state, a hallmark of the active conformation. The DFG motif (Asp-Phe-Gly) is positioned at the start of the activation loop (A-loop), acting as a critical conformational switch; its orientation (DFG-in vs. DFG-out) coordinates with the position of the C-helix to modulate the ATP-binding pocket's volume and energetic accessibility. The catalytic loop provides the necessary residues for phosphotransfer. (B) Phylogenetic tree of the human kinome[64] illustrating the breadth of the current study. Node colors categorize kinases into dark kinase domains. non-dark kinase domains and uncharacterized/non-curated domains. The model's predictive framework was validated across all major groups-including TK, TKL, STE, CK1, AGC, CAMK, and CMGC-ensuring that the PLM-derived frustration maps and binding site classifications are robust to the evolutionary divergence of the human kinome.  Kinases are obtained from the curated kinome that are visualized on the Coral kinase dendrogram.[65]  The recomputed dark kinome is shown in blue and non-dark kinases are shown in yellow. The panel was adopted from a recent study of kinome.[66]

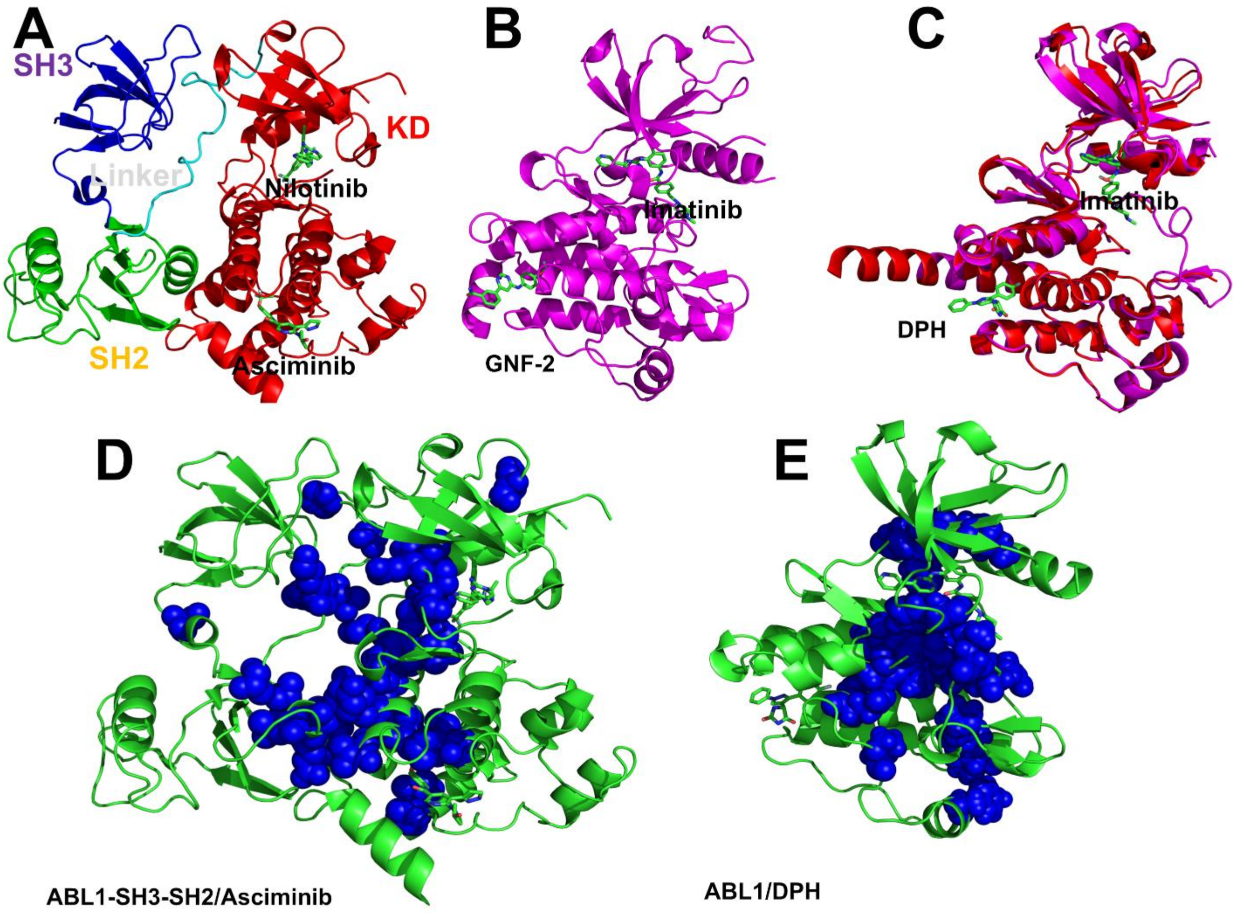


**Figure 2-figure supplement 2. Structural organization of ABL complexes with allosteric modulators and mapping of communication pathways.** (A) Architecture of the autoinhibited ABL-SH3-SH2-KD assembly. Crystal structure of the ABL regulatory core in complex with the orthosteric inhibitor Nilotinib and the allosteric inhibitor Asciminib. The multi-domain organization is indicated: SH3 domain (dark blue), SH2 domain (green), SH2-kinase linker (cyan), and the kinase domain (KD, red). Inhibitors are shown in sticks with atom-based coloring. (B) Inactive ABL-KD conformation. Structure of the isolated kinase domain (magenta) in complex with the type II inhibitor Imatinib and the allosteric myristoyl-site inhibitor GNF-2. (C) Active-like ABL-KD conformation. Structural overlay illustrating the ABL-KD in complex with Imatinib and the allosteric activator DPH. (D–E) Spatial arrangement of allosteric communication networks identified in our previous studies[67] that connect the allosteric myristoyl site and the orthosteric ATP-binding site. (D) Representative connectivity in the ABL1-SH3-SH2/Asciminib complex. (E) Preferential routes in the ABL1/DPH complex. Blue spheres represent the optimal pathways connecting the regulatory myristoyl site with the catalytic machinery.

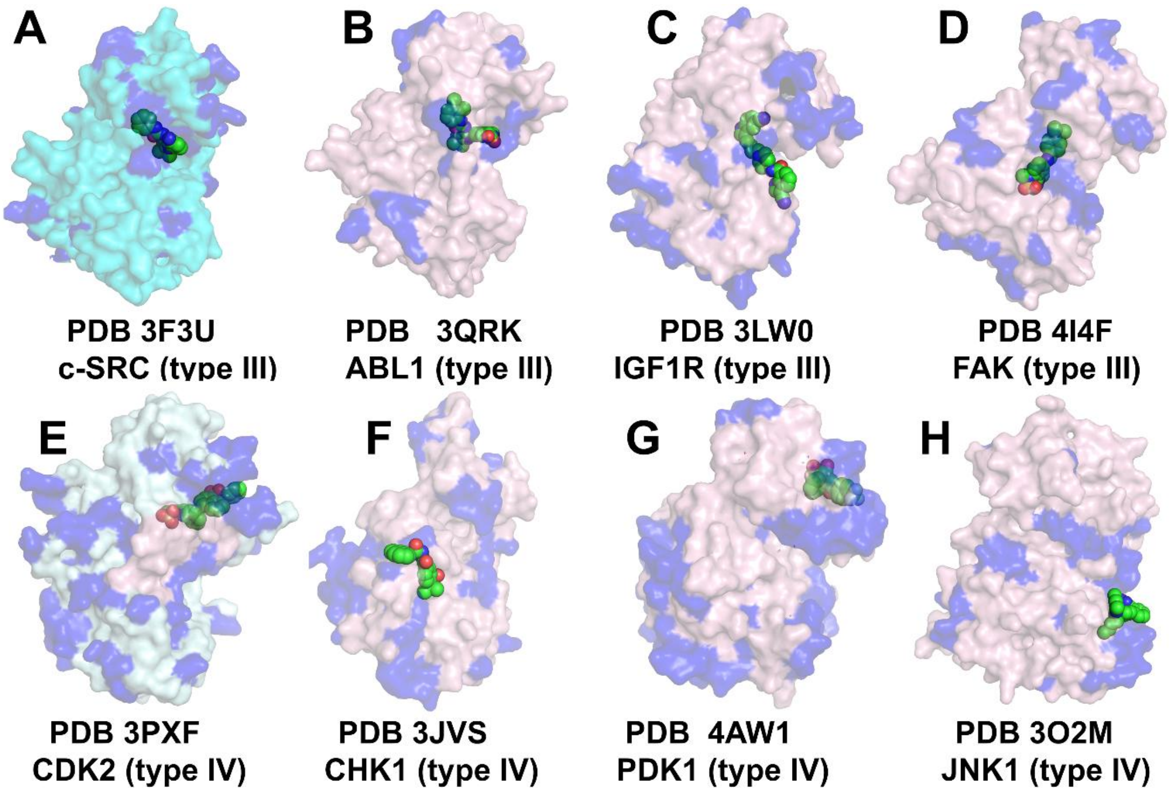


**Figure 4-figure supplement 1. Structural mapping of high neutral frustration density across diverse kinase-allosteric inhibitor complexes.** To examine the relationship between evolutionary constraints and allosteric sites, we mapped the regions of high neutral frustration density >0.7 (shown in blue) onto diverse kinase structures with type III and type IV allosteric inhibitors. Protein molecular surfaces are colored by structural origin: pale pink (Chain A) or cyan (Chain B). The corresponding allosteric inhibitors (sphere representation, atom-based coloring) are shown bound to different regulatory pockets, encompassing both Type III (adjacent to the ATP site) and Type IV (distal) modes of allosteric modulation. (Top Row) Type III adjacent-pocket inhibition. High neutral frustration density is seen localized around the orthosteric C-helix/DFG region and adjacent pockets for c-SRC (pdb 3F3U) , ABL (pdb 3QRK), IGF1R (3LW0) and FAK kinase (pdb 4I4F) (Bottom Row) Distal and Type IV allosteric inhibitors. Neutral frustration density in complexes with Type IV allosteric inhibitors CDK2 (3PXF), CHK1 (pdb 3JVS), PDK1 (4AW1) and JNK1 (pdb 3O2M). These neutrally frustrated patches illustrate the structurally diverse "allosteric blind spots."

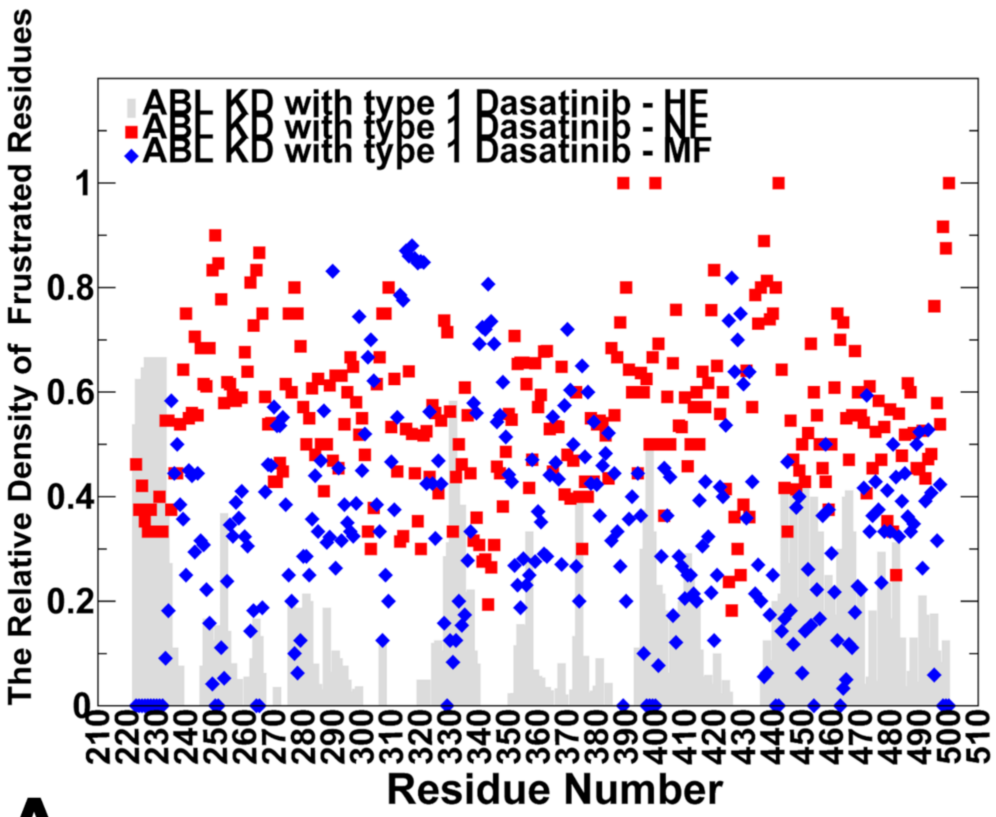
ABL KD with type 1 Dasatinib - HF
ABL KD with type 1 Dasatinib - NF
ABL KD with type 1 Dasatinib - MF
The Relative Density of Frustrated Residues
Residue Number

A

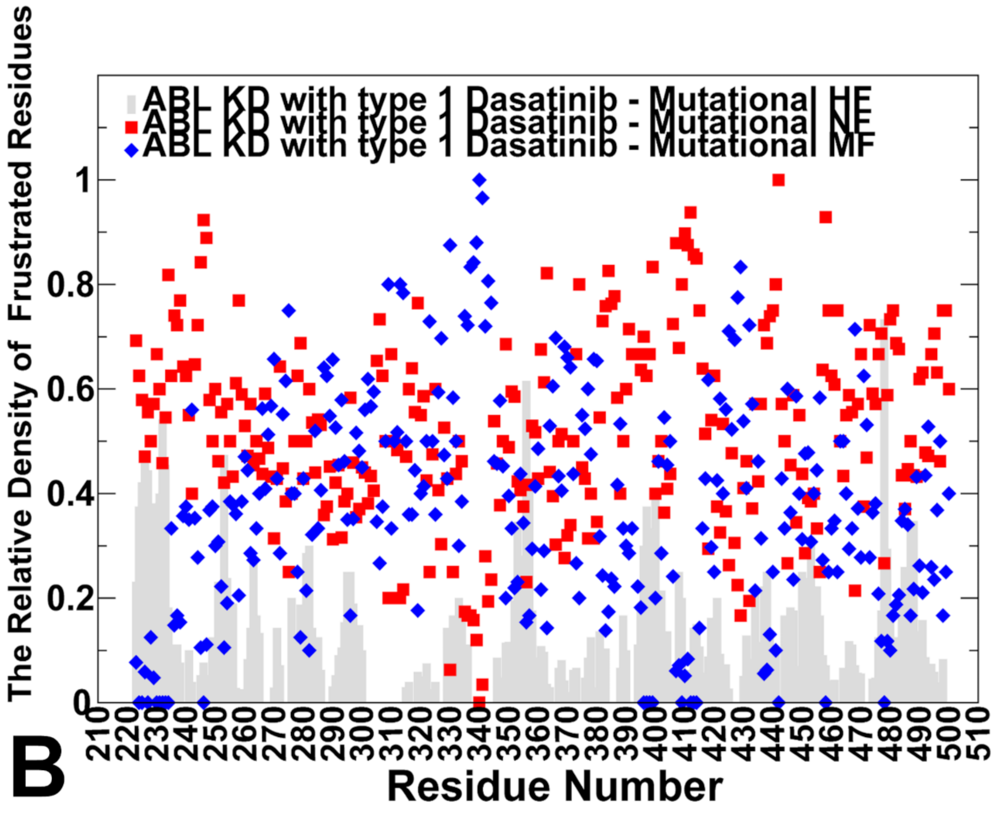
ABL KD with type 1 Dasatinib - Mutational HF
ABL KD with type 1 Dasatinib - Mutational NF
ABL KD with type 1 Dasatinib - Mutational MF
The Relative Density of Frustrated Residues
Residue Number

B

**Figure 5-figure supplements 1–5: General Legend.**

For all frustration profiles, data are categorized by frustration index: Minimally Frustrated (MF, blue diamonds), Neutrally Frustrated (NF, red squares), and Highly Frustrated (HF, grey bars). Profiles are generated using Configurational (spatial) and Mutational (evolutionary) decoy sets.

**Figure 5-figure supplement 1. ABL kinase with Dasatinib (PDB: 2GQG).**

Residue-level frustration for the kinase domain (residues 220–510) bound to a Type I inhibitor. (A) Configurational Frustration: The ATP-binding pocket and hydrophobic core exhibit high MF density, indicating structural stabilization. NF and HF residues are restricted to peripheral loops. (B) Mutational Frustration: Strong agreement between MF clusters confirms the orthosteric site is both energetically and evolutionarily optimized, establishing a "locked" baseline.

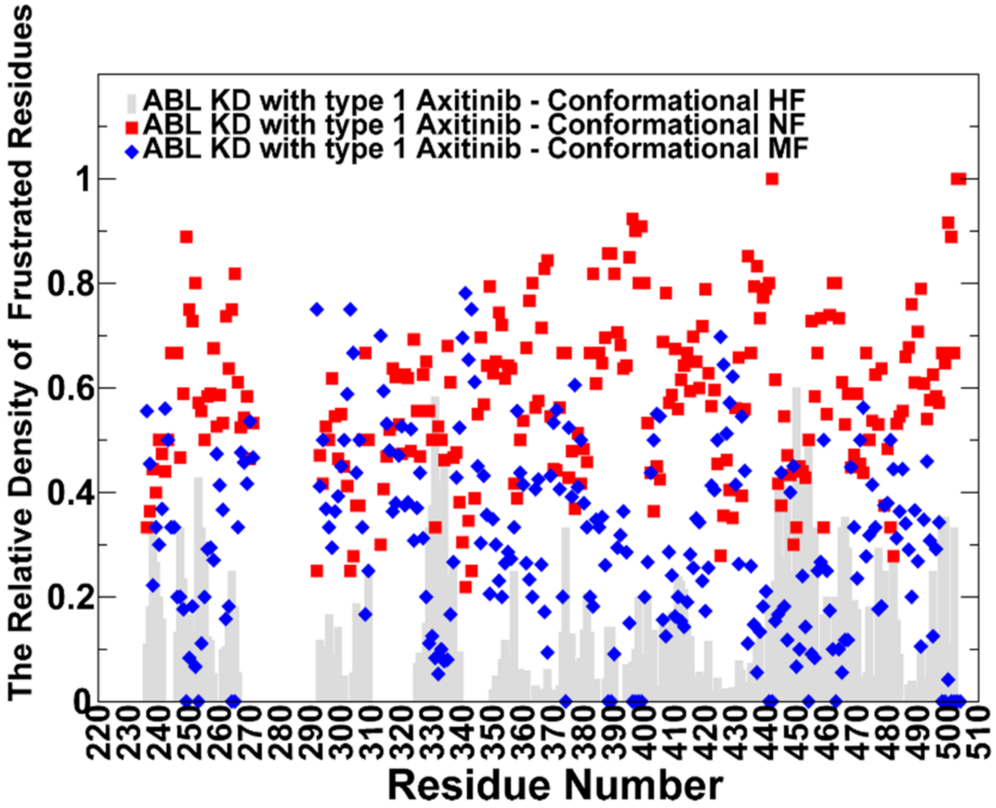
ABL KD with type 1 Axitinib - Conformational HF
ABL KD with type 1 Axitinib - Conformational NF
ABL KD with type 1 Axitinib - Conformational MF
The Relative Density of Frustrated Residues
Residue Number
0
0.2
0.4
0.6
0.8
1
220
230
240
250
260
270
280
290
300
310
320
330
340
350
360
370
380
390
400
410
420
430
440
450
460
470
480
490
500
510

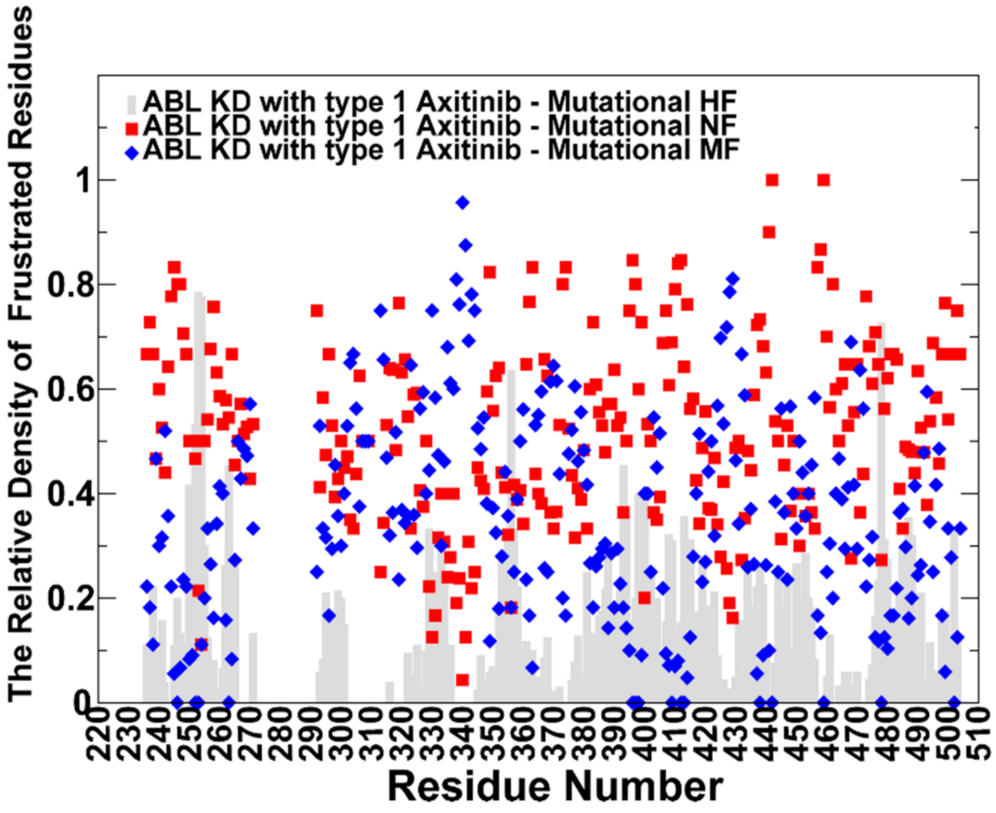
ABL KD with type 1 Axitinib - Mutational HF
ABL KD with type 1 Axitinib - Mutational NF
ABL KD with type 1 Axitinib - Mutational MF
The Relative Density of Frustrated Residues
Residue Number
0
0.2
0.4
0.6
0.8
1
220
230
240
250
260
270
280
290
300
310
320
330
340
350
360
370
380
390
400
410
420
430
440
450
460
470
480
490
500
510

**Figure 5-figure supplement 2. ABL kinase with Axitinib (PDB: 4AW9).**

Validation of orthosteric signatures using the Type I inhibitor Axitinib (residues 230–510). (A) Configurational Frustration: Dense MF interactions within the catalytic machinery reflect a stabilized active-like conformation. Gaps (residues 275–295) denote crystallographic disorder. (B) Mutational Frustration: Preservation of MF clusters confirms that the orthosteric pocket is evolutionarily optimized, providing a reference for comparison with allosteric regions.

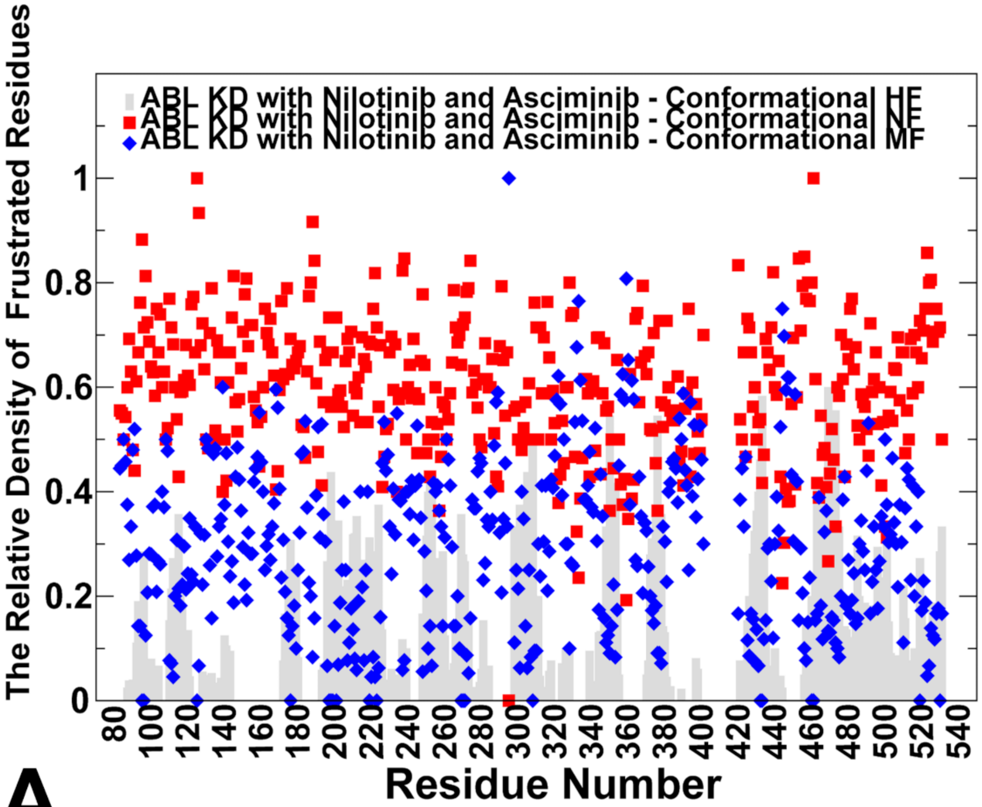

ABL KD with Nilotinib and Asciminib - Conformational HF
ABL KD with Nilotinib and Asciminib - Conformational NF
ABL KD with Nilotinib and Asciminib - Conformational MF
The Relative Density of Frustrated Residues
Residue Number


A

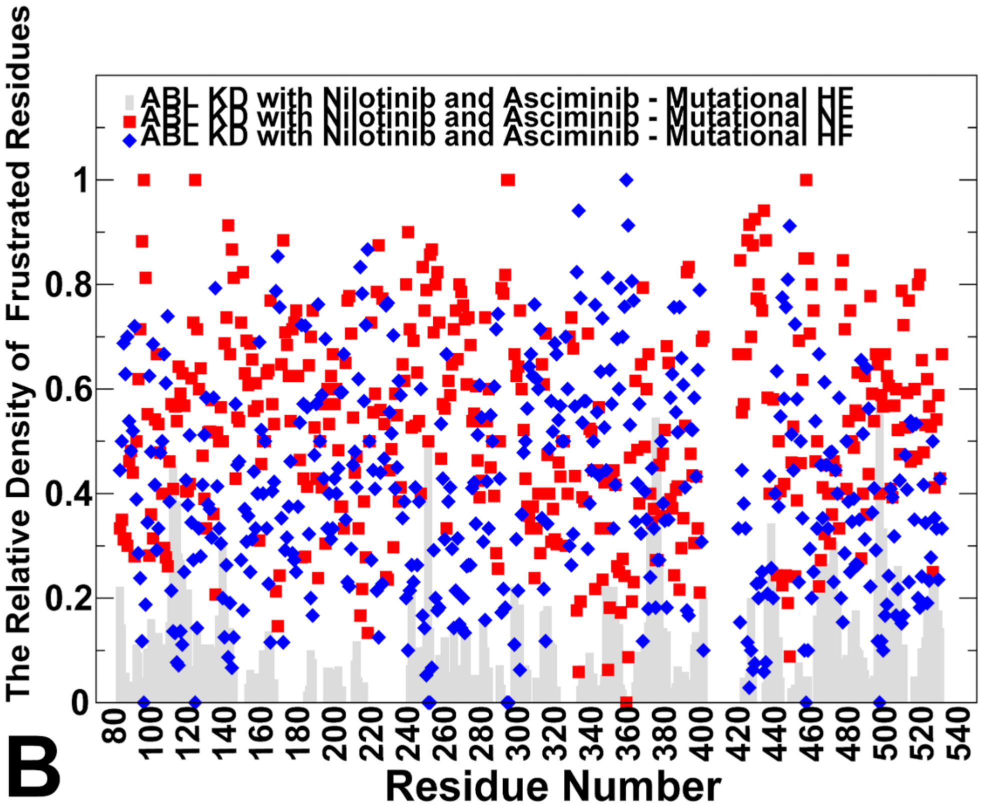

ABL KD with Nilotinib and Asciminib - Mutational HF
ABL KD with Nilotinib and Asciminib - Mutational NF
ABL KD with Nilotinib and Asciminib - Mutational HF
The Relative Density of Frustrated Residues
Residue Number


B

**Figure 5-figure supplement 3. Autoinhibited ABL-SH3-SH2-KD complex (PDB: 5MO4).**

Analysis of the full regulatory assembly bound to Nilotinib (orthosteric) and Asciminib (allosteric). (A) Configurational Frustration: Widespread MF clusters define the domain interfaces of this "clamped" state (residues 80–540). Notably, the myristoyl pocket remains dominated by NF residues despite Asciminib occupancy. (B) Mutational Frustration: Persistent NF density confirms that the allosteric site maintains a neutral energetic signature independent of ligand-induced stabilization.

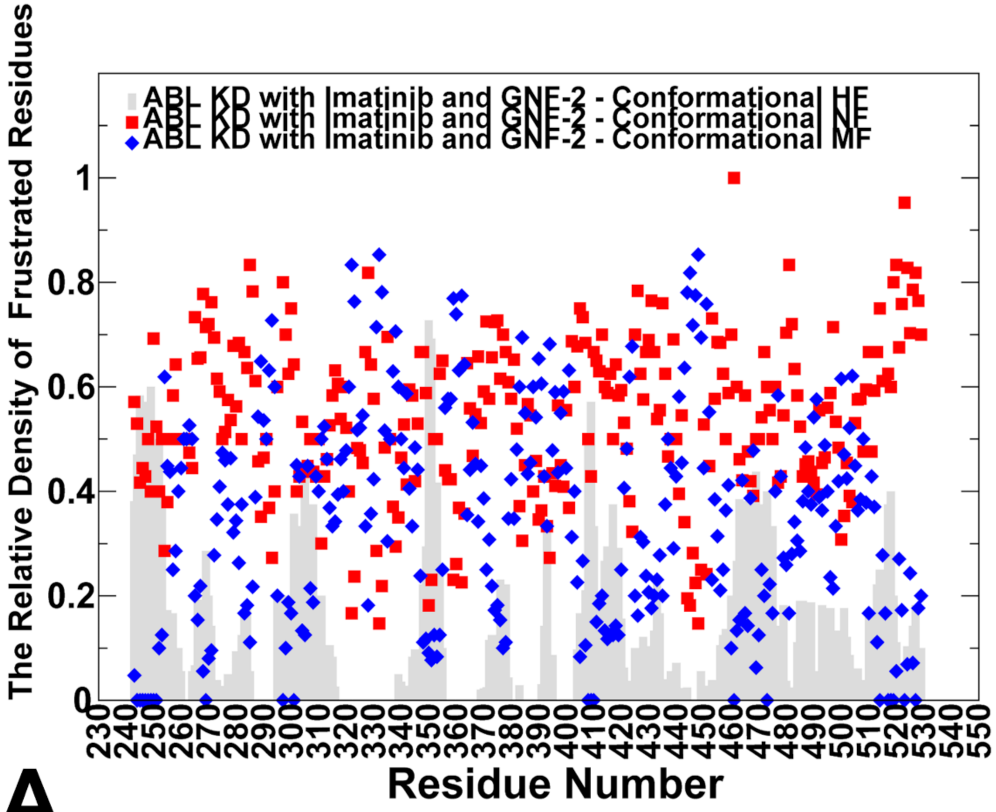
ABL KD with Imatinib and GNF-2 - Conformational HF
ABL KD with Imatinib and GNF-2 - Conformational NF
ABL KD with Imatinib and GNF-2 - Conformational MF
The Relative Density of Frustrated Residues
Residue Number
0
0.2
0.4
0.6
0.8
1
230 240 250 260 270 280 290 300 310 320 330 340 350 360 370 380 390 400 410 420 430 440 450 460 470 480 490 500 510 520 530 540 550

A

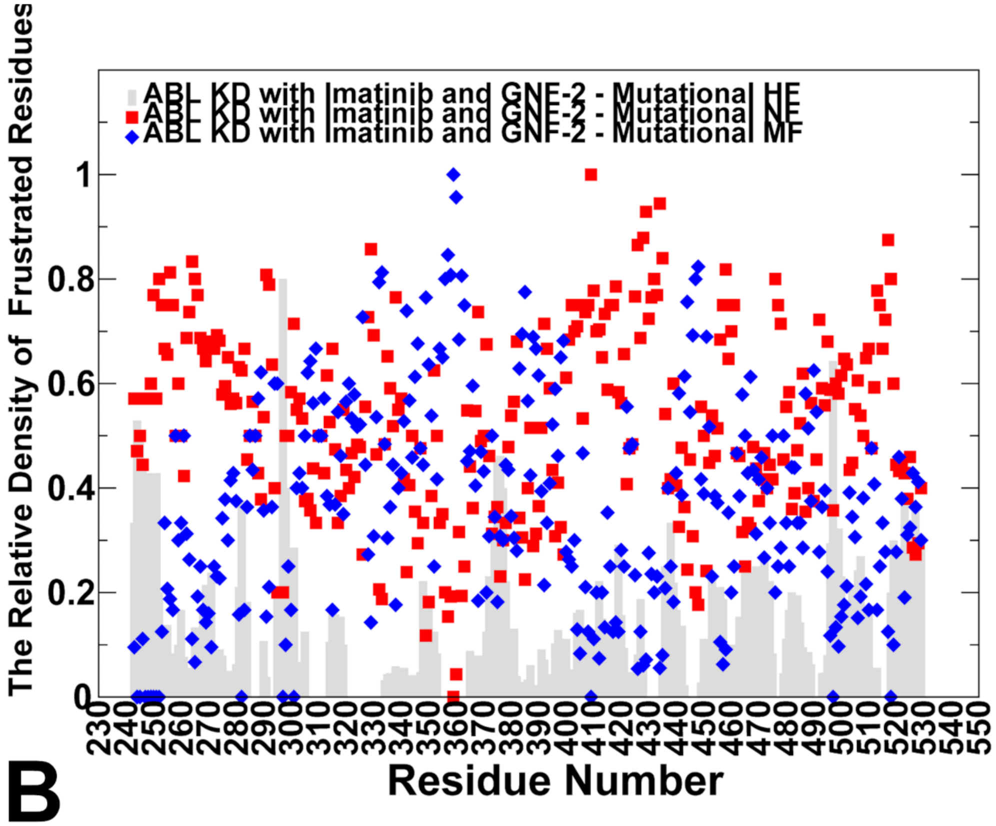
ABL KD with Imatinib and GNF-2 - Mutational HF
ABL KD with Imatinib and GNF-2 - Mutational NF
ABL KD with Imatinib and GNF-2 - Mutational MF
The Relative Density of Frustrated Residues
Residue Number
0
0.2
0.4
0.6
0.8
1
230 240 250 260 270 280 290 300 310 320 330 340 350 360 370 380 390 400 410 420 430 440 450 460 470 480 490 500 510 520 530 540 550

B

**Figure 5-figure supplement 4. ABL kinase with Imatinib and GNF-2 (PDB: 3K5V).**

Analysis of the kinase domain (residues 230–530) with dual occupancy of the ATP (Imatinib) and myristoyl (GNF-2) pockets. (A) Configurational Frustration: The dual inhibitors stabilize the inactive DFG-out conformation. While the core shows significant MF clusters, the allosteric myristoyl pocket and surrounding network maintain high NF density. (B) Mutational Frustration: Persistent NF density confirms the lack of rigid evolutionary optimization at the allosteric interface.

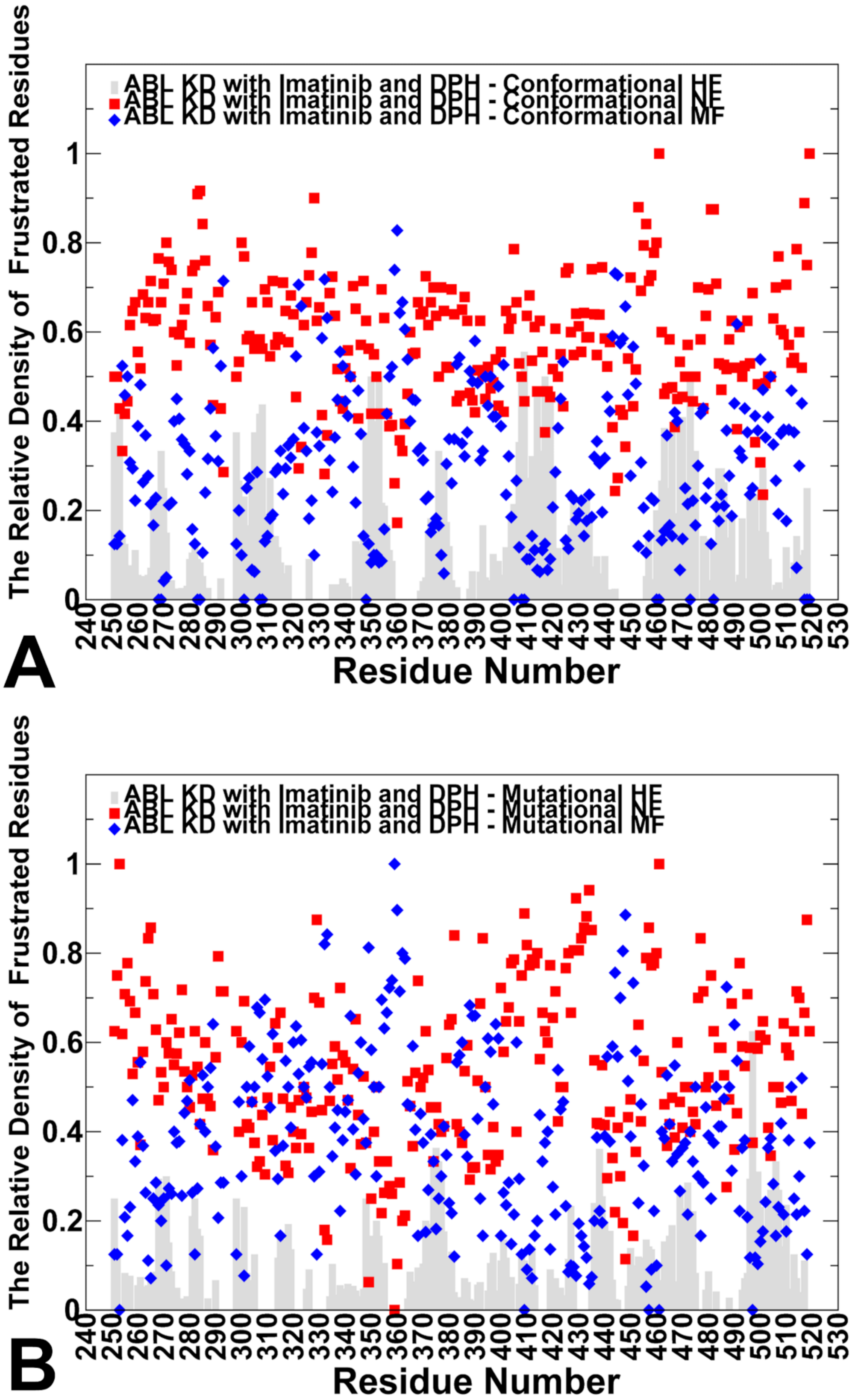

ABL KD with Imatinib and DPH - Conformational HF
ABL KD with Imatinib and DPH - Conformational NF
ABL KD with Imatinib and DPH - Conformational MF
The Relative Density of Frustrated Residues
Residue Number
A
ABL KD with Imatinib and DPH - Mutational HF
ABL KD with Imatinib and DPH - Mutational NF
ABL KD with Imatinib and DPH - Mutational MF
The Relative Density of Frustrated Residues
Residue Number
B

**Figure 5-figure supplement 5. ABL kinase with Imatinib and the activator DPH (PDB: 3PYY).**

Analysis of the kinase domain (residues 240–530) bound to an allosteric activator. (A) Configurational Frustration: Unlike allosteric inhibitors, the activator DPH results in a heterogeneous landscape. While the orthosteric pocket remains stable (MF), the allosteric site exhibits high NF and HF density, reflecting the energetic strain required to disrupt autoinhibition. (B) Mutational Frustration: The allosteric machinery maintains high NF density even under activation, highlighting the shared neutral signature of regulatory pockets.
