## Supplementary figures and images for "Decoding Allosteric Grammar with Explainable AI Integrating Protein Language Models and Energy Landscape Analysis: Neutral Frustration at Allosteric Binding Sites Encodes Regulatory Versatility in Protein Kinases"

### Figure 1-figure supplement 1.tif

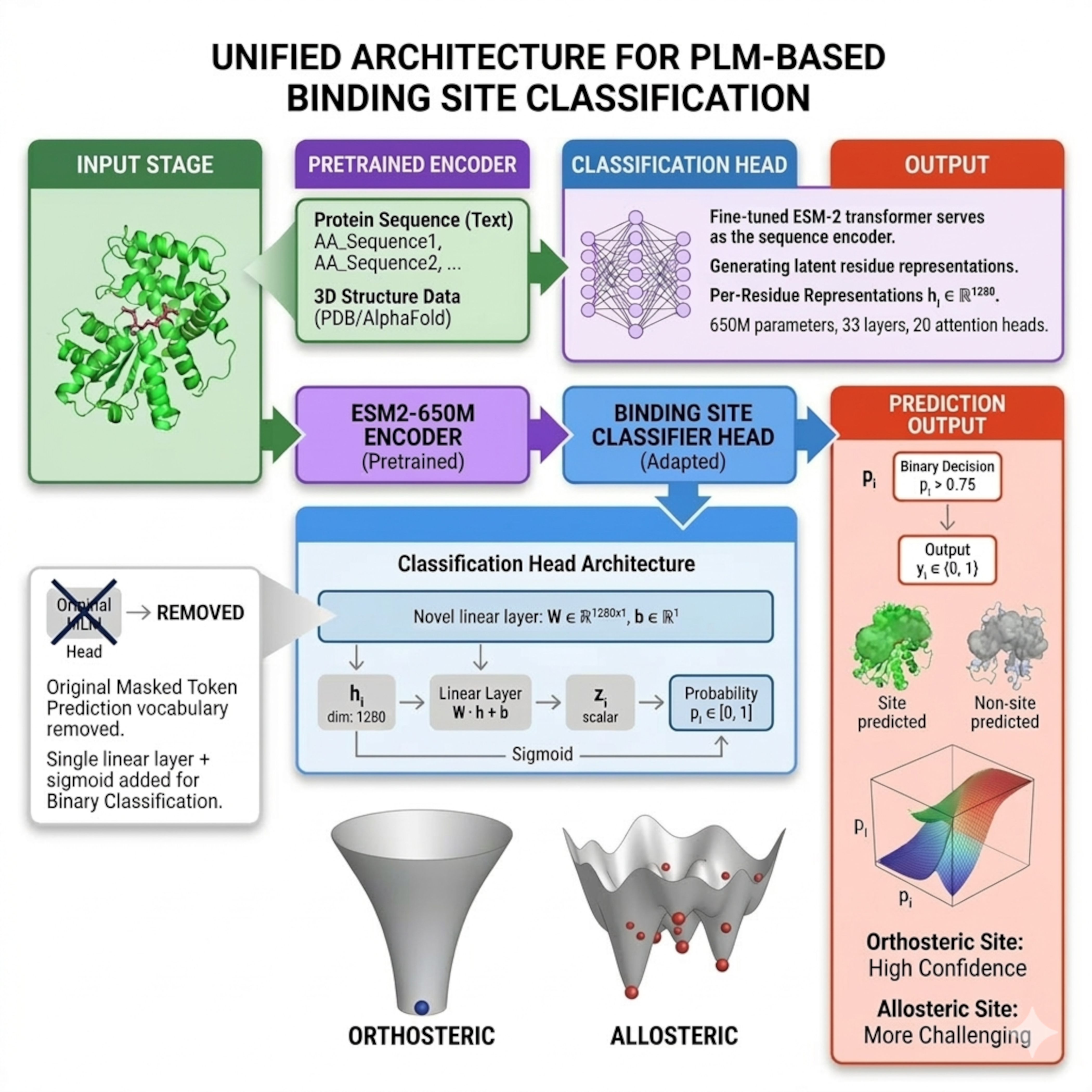

### Figure 2-figure supplement 1.tif

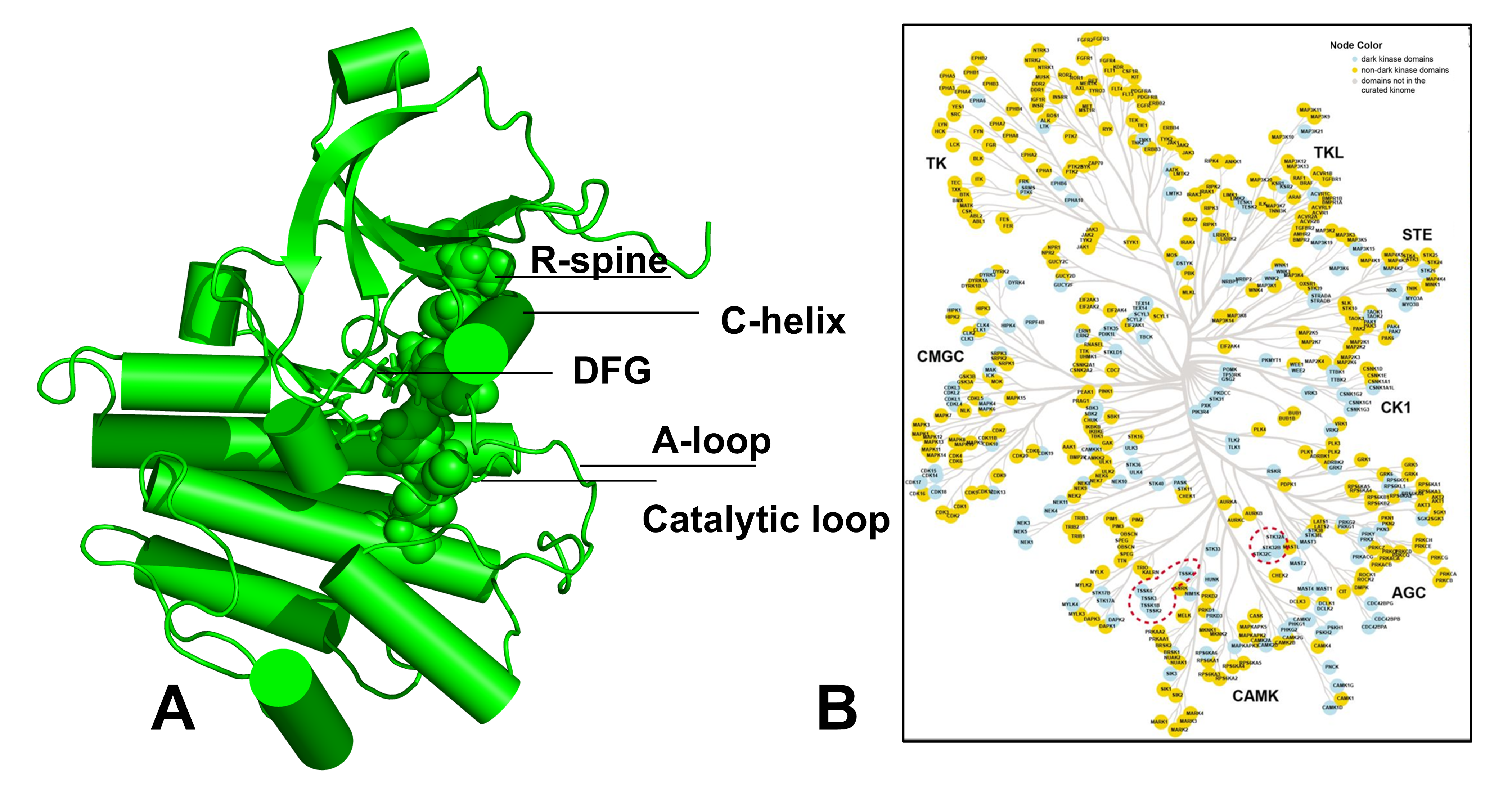

### Figure 2-figure supplement 2.tif

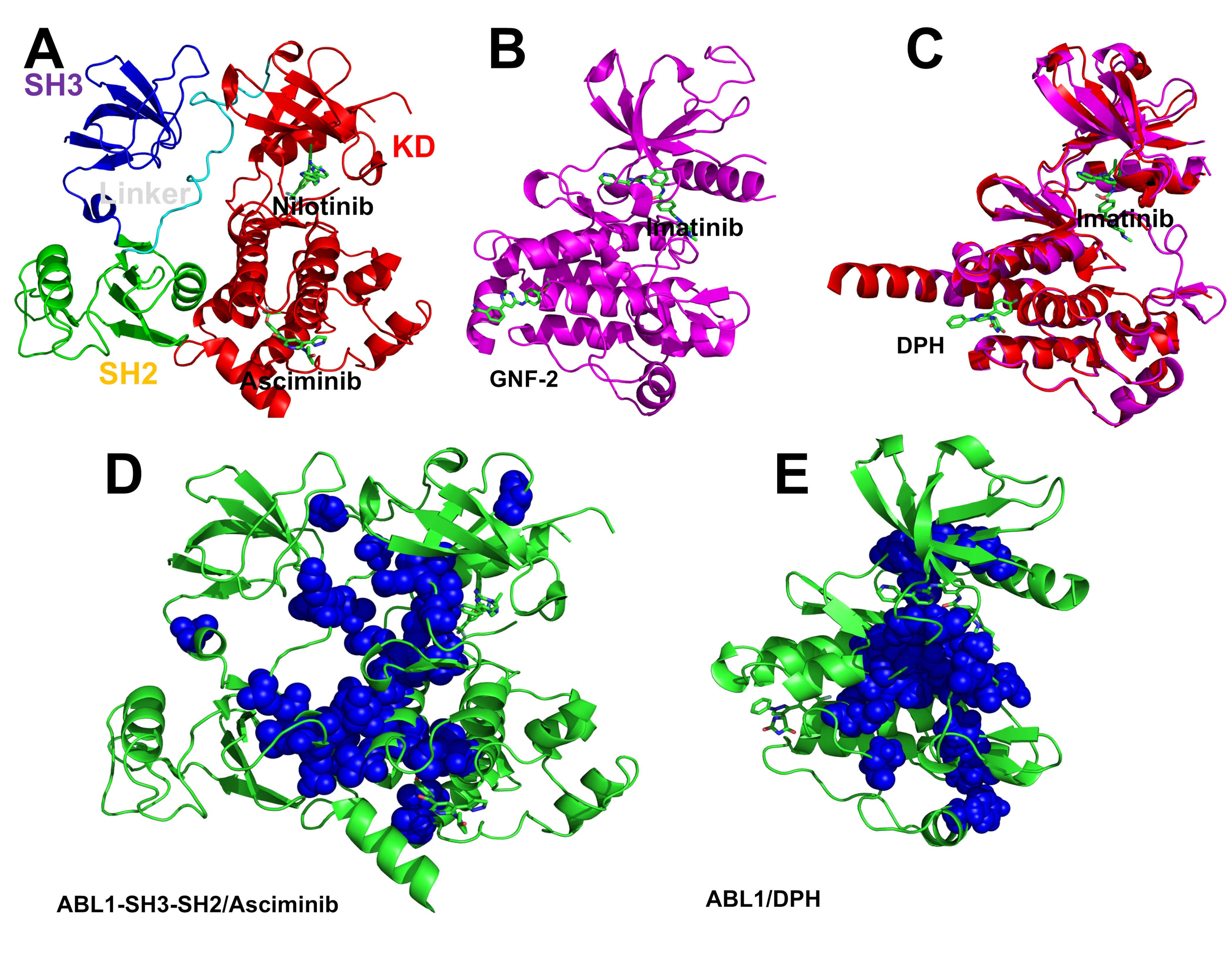

### Figure 4-figure supplement 1.tif

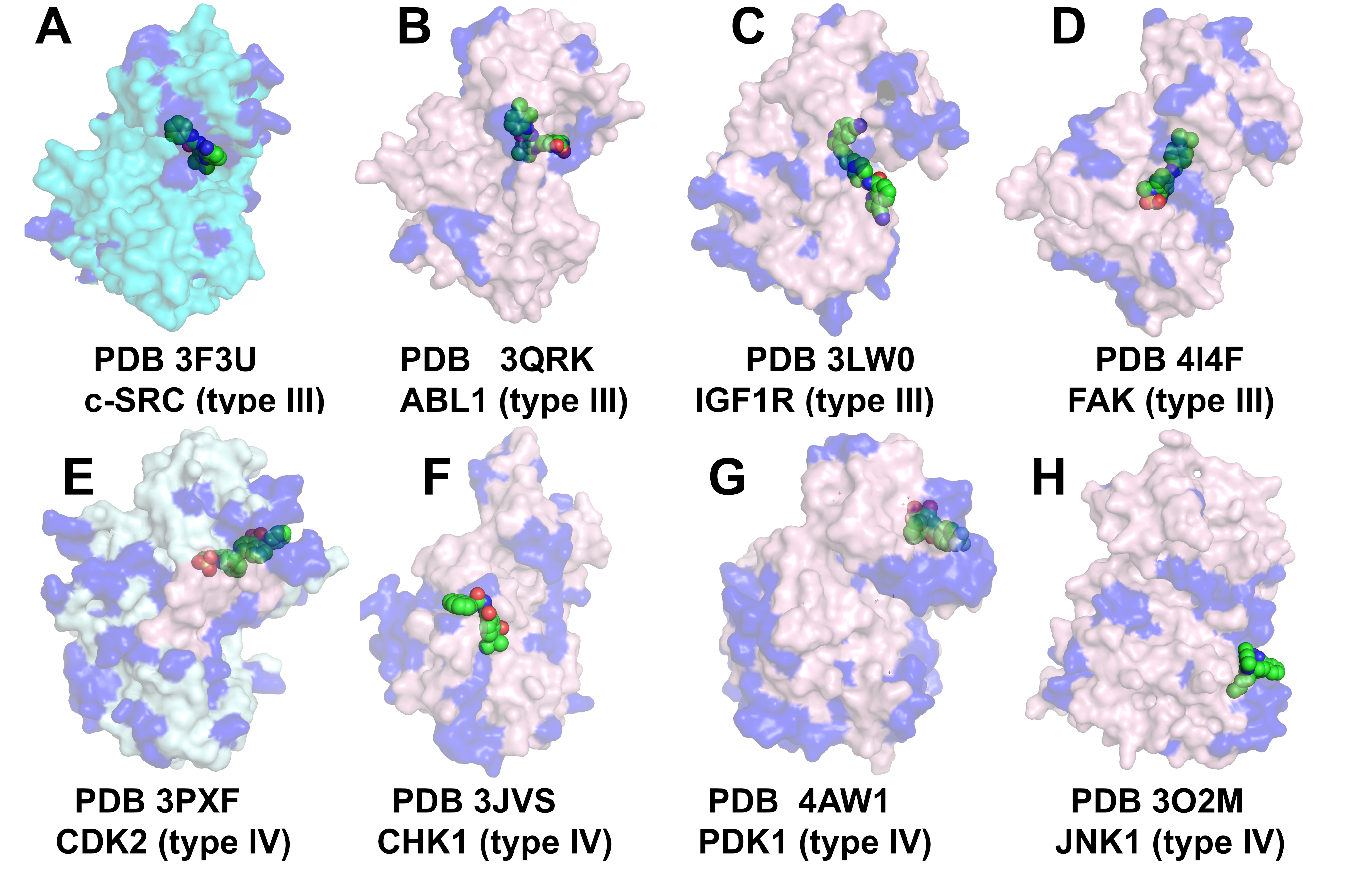

### Figure 5-figure supplement 1.tif

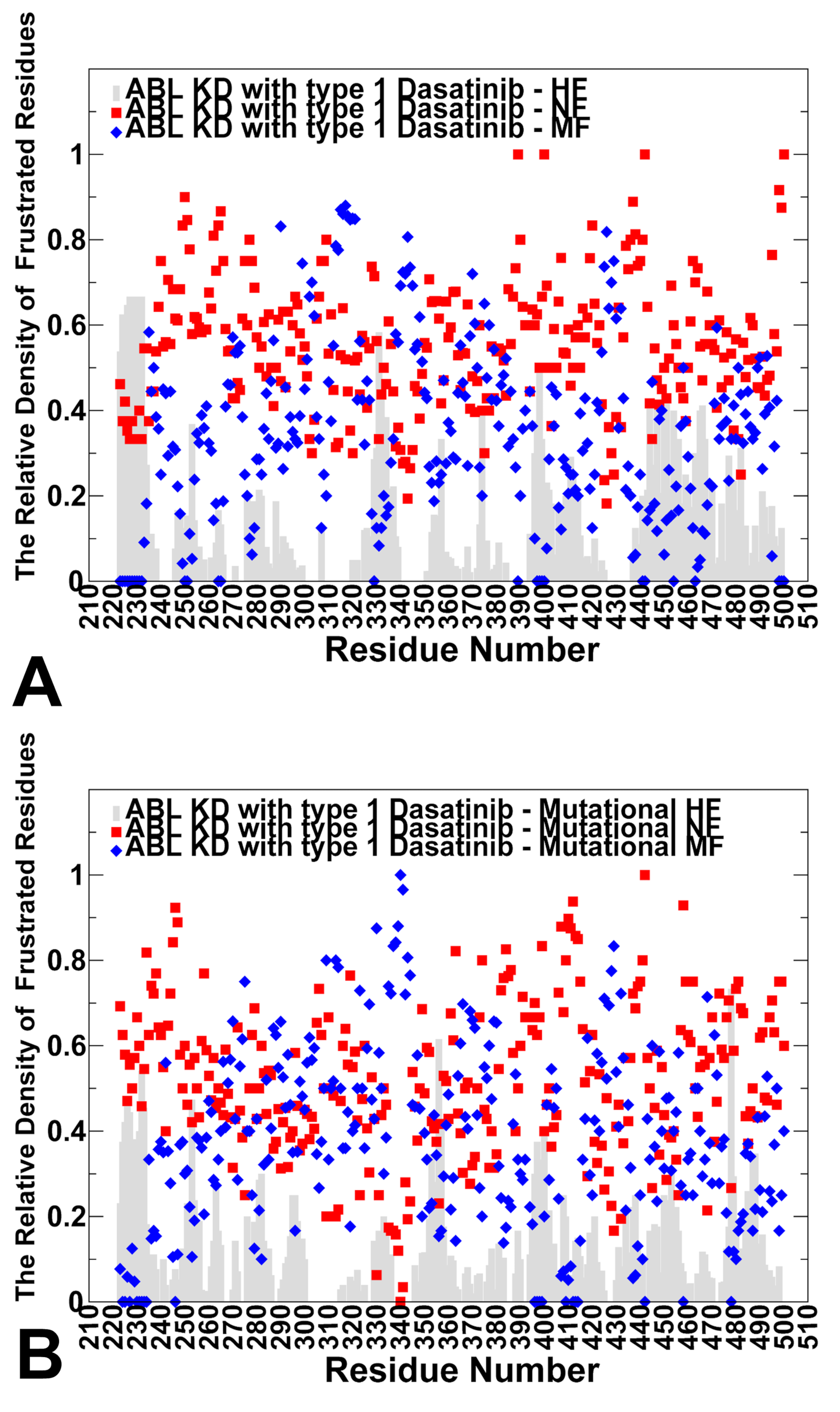

### Figure 5-figure supplement 2.tif

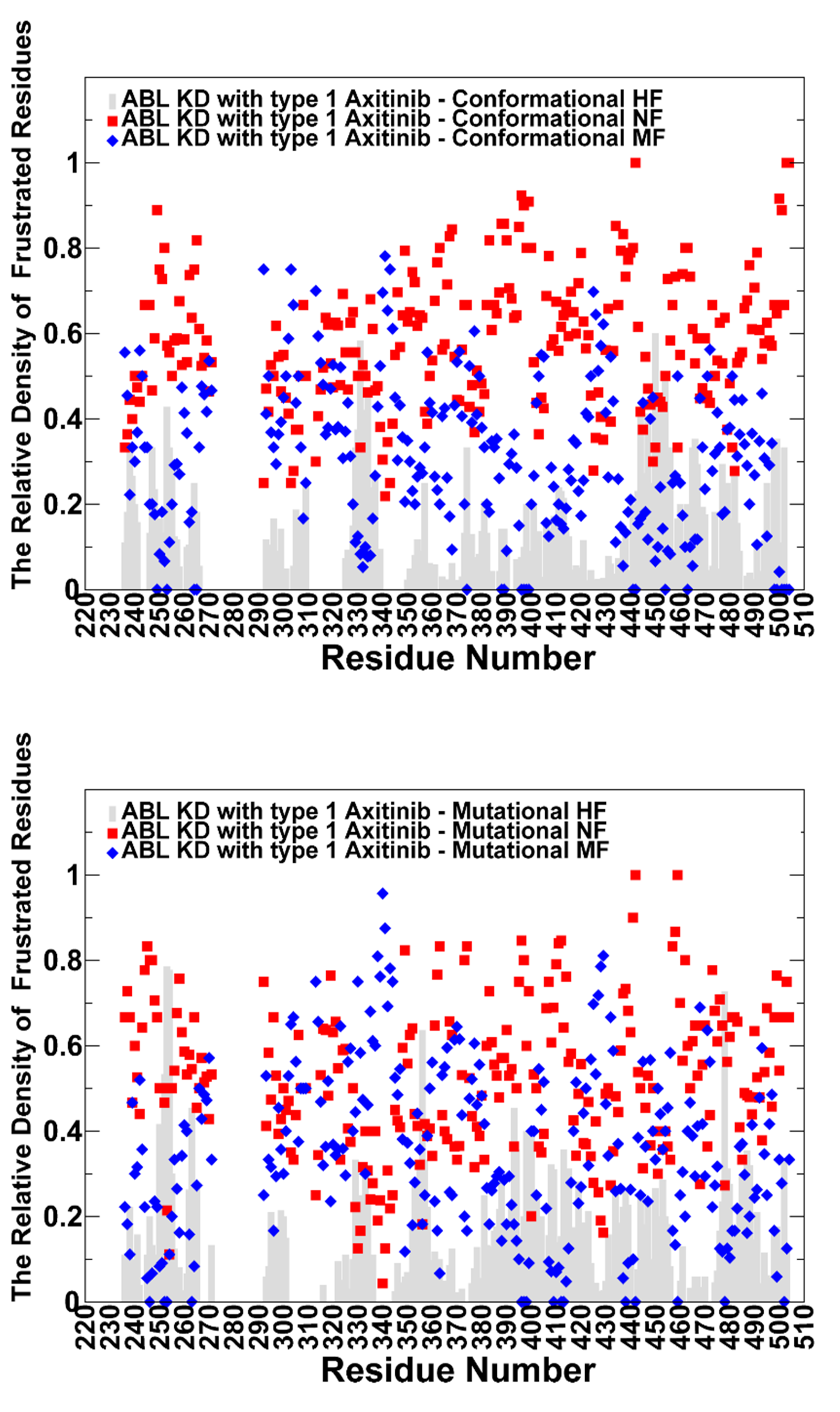

### Figure 5-figure supplement.3.tif

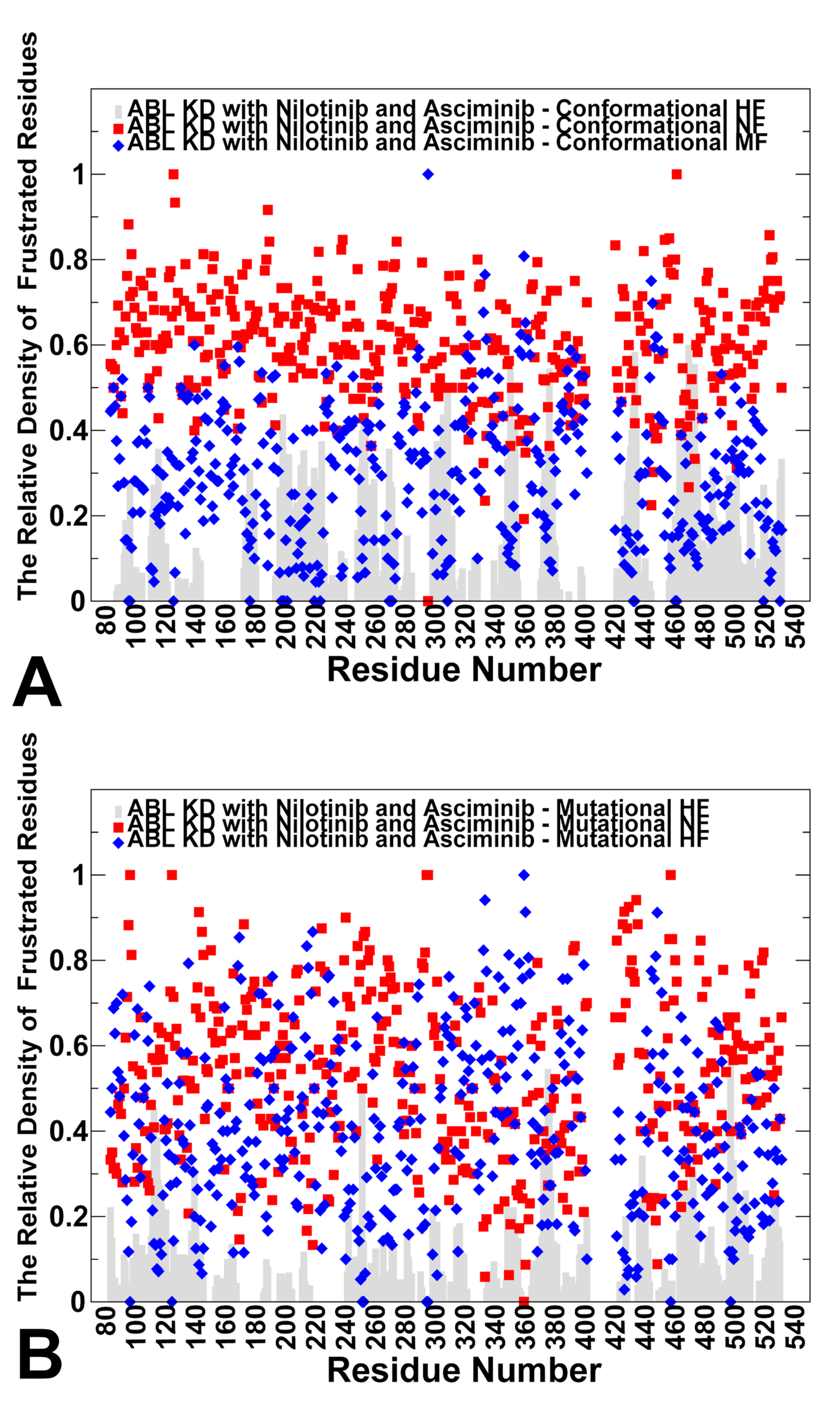

### Figure 5-figure supplement.4.tif

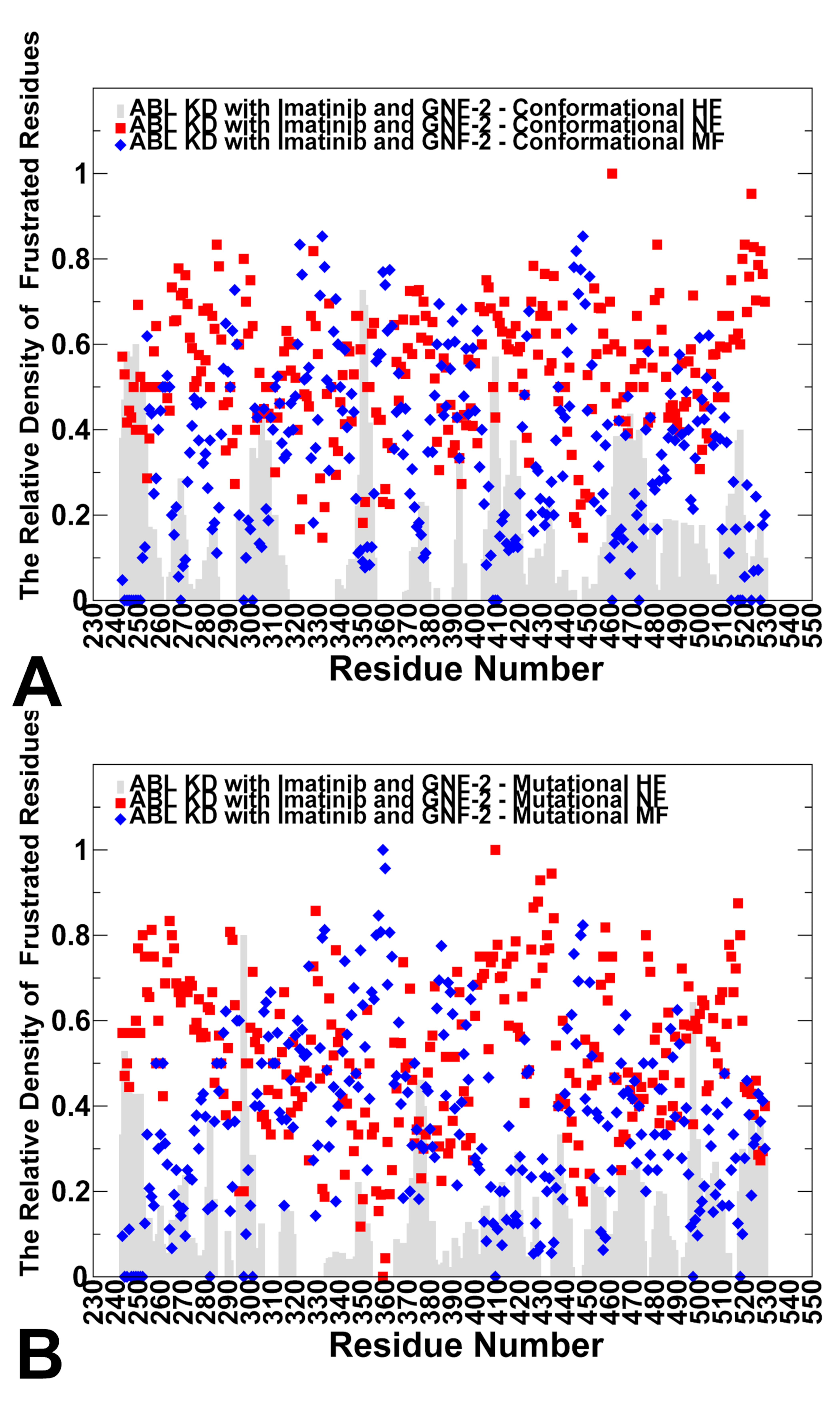

### Figure 5-figure supplement.5.tif

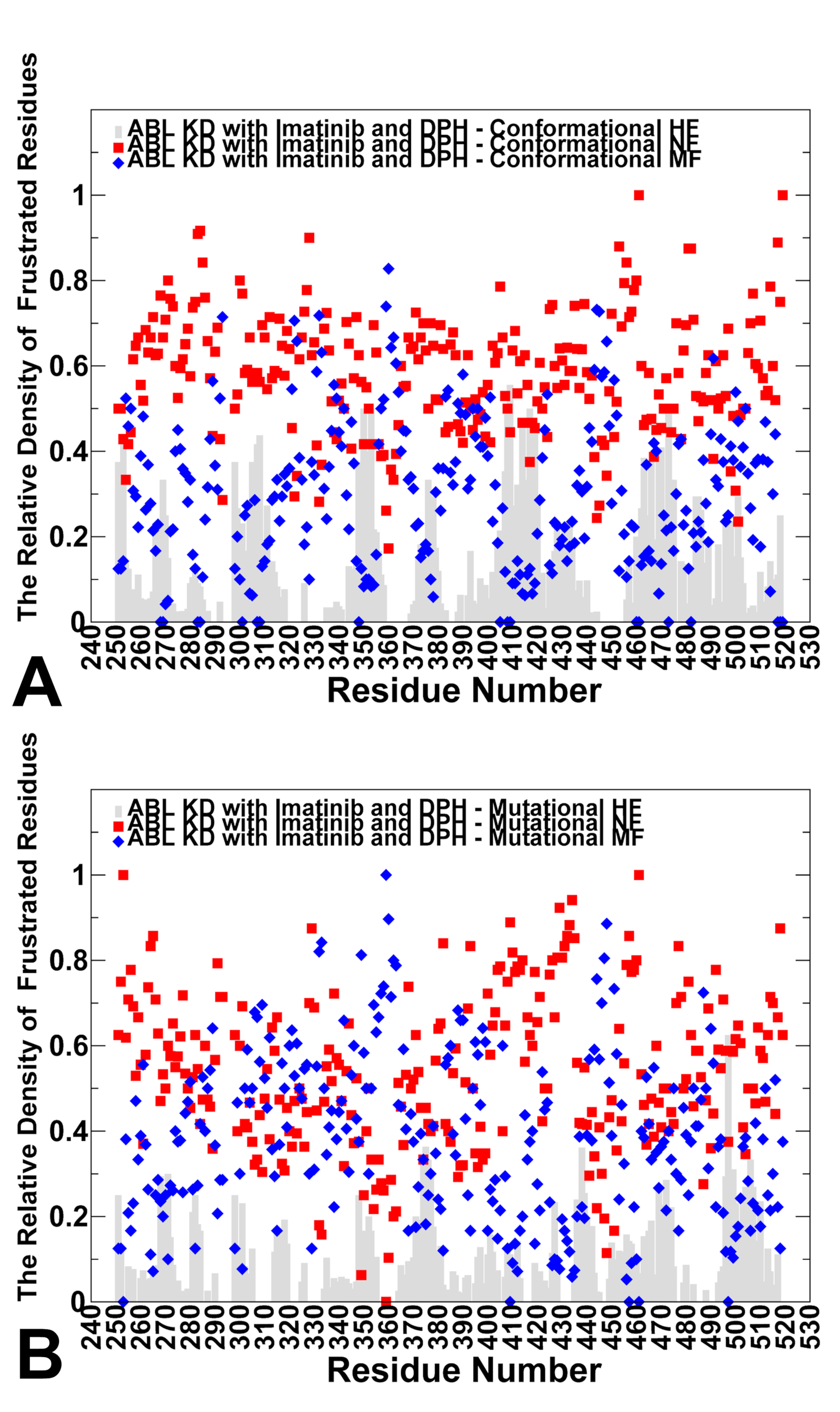
